# Ultrapotent bispecific antibodies neutralize emerging SARS-CoV-2 variants

**DOI:** 10.1101/2021.04.01.437942

**Authors:** Hyeseon Cho, Kristina Kay Gonzales-Wartz, Deli Huang, Meng Yuan, Mary Peterson, Janie Liang, Nathan Beutler, Jonathan L. Torres, Yu Cong, Elena Postnikova, Sandhya Bangaru, Chloe Adrienna Talana, Wei Shi, Eun Sung Yang, Yi Zhang, Kwanyee Leung, Lingshu Wang, Linghang Peng, Jeff Skinner, Shanping Li, Nicholas C. Wu, Hejun Liu, Cherrelle Dacon, Thomas Moyer, Melanie Cohen, Ming Zhao, F. Eun-Hyung Lee, Rona S. Weinberg, Iyadh Douagi, Robin Gross, Connie Schmaljohn, Amarendra Pegu, John R. Mascola, Michael Holbrook, David Nemazee, Thomas F. Rogers, Andrew B. Ward, Ian A. Wilson, Peter D. Crompton, Joshua Tan

## Abstract

The emergence of SARS-CoV-2 variants that threaten the efficacy of existing vaccines and therapeutic antibodies underscores the urgent need for new antibody-based tools that potently neutralize variants by targeting multiple sites of the spike protein. We isolated 216 monoclonal antibodies targeting SARS-CoV-2 from plasmablasts and memory B cells of COVID-19 patients. The three most potent antibodies targeted distinct regions of the RBD, and all three neutralized the SARS-CoV-2 variants B.1.1.7 and B.1.351. The crystal structure of the most potent antibody, CV503, revealed that it binds to the ridge region of SARS-CoV-2 RBD, competes with the ACE2 receptor, and has limited contact with key variant residues K417, E484 and N501. We designed bispecific antibodies by combining non-overlapping specificities and identified five ultrapotent bispecific antibodies that inhibit authentic SARS-CoV-2 infection at concentrations of <1 ng/mL. Through a novel mode of action three bispecific antibodies cross-linked adjacent spike proteins using dual NTD/RBD specificities. One bispecific antibody was >100-fold more potent than a cocktail of its parent monoclonals *in vitro* and prevented clinical disease in a hamster model at a 2.5 mg/kg dose. Notably, six of nine bispecific antibodies neutralized B.1.1.7, B.1.351 and the wild-type virus with comparable potency, despite partial or complete loss of activity of at least one parent monoclonal antibody against B.1.351. Furthermore, a bispecific antibody that neutralized B.1.351 protected against SARS-CoV-2 expressing the crucial E484K mutation in the hamster model. Thus, bispecific antibodies represent a promising next-generation countermeasure against SARS-CoV-2 variants of concern.

## Introduction

The novel coronavirus SARS-CoV-2 emerged in Wuhan, China in 2019 and has infected 129 million individuals and caused 2.8 million deaths worldwide at the time of writing. *In vitro* and *in vivo* experiments, along with observational studies in humans, strongly support a role for SARS-CoV-2-neutralizing antibodies in protection against COVID-19, but emerging SARS-CoV-2 variants such as B.1.1.7 (UK), B.1.351 (South Africa) and P.1 (Brazil) harbor mutations that may decrease the efficacy of existing vaccines and therapeutic monoclonal antibodies (mAbs), underscoring the importance of developing new antibody-based tools that potently neutralize variants by targeting diverse sites of the spike protein^1–17^. To date, most studies investigating SARS-CoV-2-specific mAbs have used antigen probe-based methods to isolate memory B cells (MBCs) or a mixture of plasmablasts and MBCs^11, 12, 18–21^. Here we used an approach that does not rely on antigen probes to generate a large panel of mAbs from both plasmablasts and MBCs of recovered COVID-19 patients. We combined potent monoclonal antibodies with non-overlapping specificities to generate bispecific antibodies targeting multiple regions of the spike protein that potently neutralize emerging SARS-CoV-2 variants.

### Characterization of plasma from COVID-19 convalescent donors

To decipher the characteristics of circulating antibodies in individuals who successfully controlled SARS-CoV-2 infection, we first examined convalescent plasma samples of 126 individuals in New York City who had recovered from PCR-documented SARS-CoV-2 infection. Samples were collected in April 2020 and therefore reflect the B cell response during the first outbreak in the study area. We tested plasma for binding to the spike protein of non-SARS-CoV-2 coronaviruses, as well as to the receptor-binding domain (RBD) and N-terminal domain (NTD) of SARS-CoV-2 (Supp. Fig. 1A, see Methods for details of antigen production). All subjects had detectable levels of antibodies to at least one non-SARS-CoV-2 spike protein, consistent with previous exposure to seasonal coronaviruses. As expected, most subjects also had detectable antibodies to the SARS-CoV-2 spike protein (119/126), RBD (106/126) and NTD (122/126), and antibody levels against these targets correlated with each other (Supp. Fig. 1B). We then tested plasma for neutralization of authentic wild-type SARS-CoV-2 and found a wide range of neutralizing titers from <40 to 765 (Supp. Fig. 1A). Neutralization potency weakly correlated with antibody levels to SARS-CoV-2 spike, RBD and NTD, and several plasma samples were non-neutralizing despite high antibody levels to each of these targets (Supp. Fig. 1C), suggesting that the fine specificity of the antibody epitopes is critical for effective neutralization of SARS-CoV-2.

**Fig. 1.**
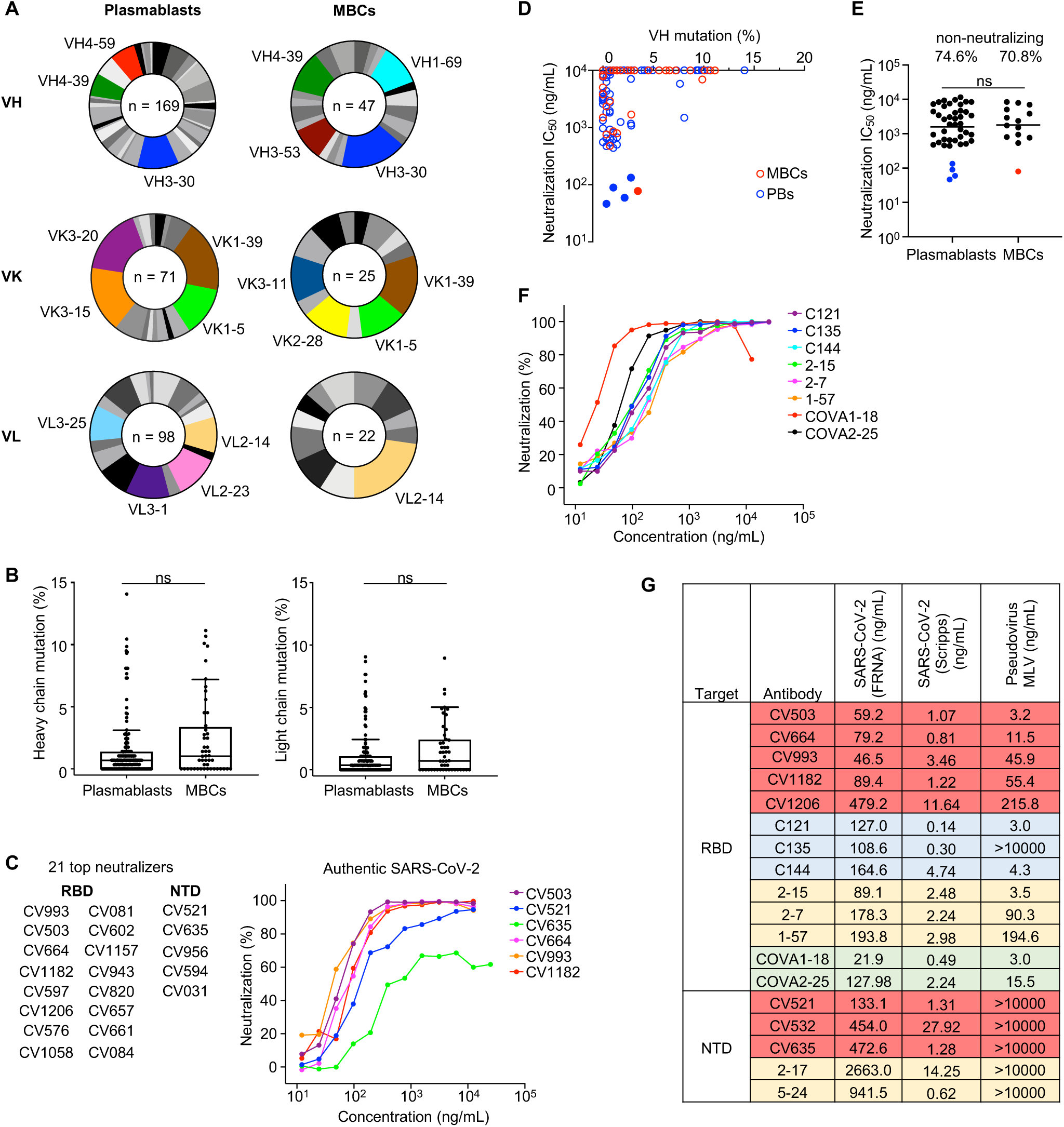
Plasmablasts produce potent antibodies against SARS-CoV-2 with diverse V genes and limited mutations. **A,** VH, VK and VL gene usage of antibodies from plasmablasts and memory B cells (MBCs). Up to the top four genes in each chart are shown with different colors (genes that were tied for 4^th^ and lower are not highlighted). **B**, Tukey’s plots showing heavy and light chain gene mutations of antibodies from plasmablasts and MBCs. Percent mutations were compared with the Mann-Whitney U-test. The middle line shows the median, and the box extends from the 1^st^ to 3^rd^ quartile. **C**, Top 21 neutralizing antibodies (IC50 <1 μg/mL) by antigen specificity (left), and neutralization curves of selected antibodies (right). **D**, SARS-CoV-2 neutralization potency versus heavy chain mutation levels of antibody panel. The 5 most potent antibodies are shown as solid circles. **E**, Neutralization potency of antibodies by cell type. Top values indicate percentages of non-neutralizing antibodies. Horizontal bars indicate mean values; Mann-Whitney U-test (non-neutralizing antibodies excluded from calculation). The 5 most potent antibodies are shown in color. **F**, SARS-CoV-2 neutralization curves of benchmark antibodies from different groups^11, 18, 21^. **G,** Neutralization IC50 values of antibodies in our panel and benchmark antibodies in three different neutralization assays. Authentic SARS-CoV-2 FRNA values are from a single experiment done in quadruplicate, authentic SARS-CoV-2 (Scripps) values are an average of two experiments done in duplicate, pseudovirus (MLV) values are an average of 3 experiments in duplicate. The colors indicate the different sources of the antibodies: red, this study; blue, ref. 21; yellow, ref. 11; green, ref. 18.

### SARS-CoV-2-specific antibodies derived from plasmablasts and memory B cells use diverse V genes and have few mutations

The specificity and potency of mAbs derived from SARS-CoV-2-specific plasmablasts is poorly characterized^22, 23^. Therefore, we developed a single-cell assay using the Berkeley Lights Beacon optofluidics device to screen for SARS-CoV-2-specific mAbs secreted by plasmablasts *ex vivo* as well as MBCs following *in vitro* stimulation. Of the 126 donors, we focused on the nine whose plasma most potently neutralized SARS-CoV-2 *in vitro,* as well as three moderate/poor neutralizers as comparators (Supp. Fig. 1A). Circulating plasmablasts (CD19^+^CD27^++^CD38^++^) from these donors were bulk FACS-sorted, distributed individually into nanoliter-volume pens in a microfluidics chip, and then screened directly for secretion of antibodies that bound to beads coated with SARS-CoV-2 spike or RBD (Supp. Fig. 1D). A total of ∼44,000 plasmablasts were screened using this assay, of which 787 supernatants bound to spike and/or RBD. In parallel, we obtained 291 positive supernatants from MBCs (CD19^+^IgG^+^/IgA^+^) that had been activated *in vitro* and screened in the same assay. We exported B cells of interest from the microfluidics chip, performed reverse-transcription PCR to obtain heavy and light chain sequences, and expressed the antibodies recombinantly. In total, we expressed and characterized 169 mAbs targeting SARS-CoV-2 from plasmablasts and 47 from MBCs from the same 12 individuals (Supp. Fig. 1E, Supplementary Table 1). Of the plasmablast-derived mAbs, 59 targeted the RBD, 64 targeted the NTD, and 46 targeted neither (S2-specific or possibly quaternary), indicating a response to SARS-CoV-2 that is distributed along the entire spike protein. Antibodies isolated from both plasmablasts and MBCs used diverse V genes, with many of the enriched gene families matching those previously reported^18, 24^ (Fig. 1A) and partially overlapping between plasmablasts and MBCs. For instance, while genes such as *VH3-30* and *VH4-39* were enriched in both groups, *VH3-53* was more common among MBCs (4^th^ most frequent) than plasmablasts (14^th^ most frequent). We also found that plasmablasts and MBCs had similarly low mutation levels (<3%) in their heavy and light chain genes (Fig. 1B), consistent with their differentiation from naïve B cells without extensive germinal center experience^11, 12, 18, 21, 25^. The markers used for cell sorting did not allow us to distinguish between activated and resting MBCs, but MBC-derived mAbs rarely recognized spike proteins of previously circulating betacoronaviruses, providing further evidence that resting MBCs were not the source of most of the isolated mAbs. A minority of mAbs from plasmablasts and MBCs did have higher mutation levels (∼10%), suggesting that they arose from pre-existing MBCs. However, even these antibodies did not cross-react with seasonal coronaviruses (Supplementary Table 1), suggesting the possibility that they may target other pathogens or even self-antigens, as recently described^26^. To minimize the effects of inter-donor variation, we analyzed mAbs isolated from donor COV050, the only individual from whom we obtained similar numbers of plasmablast- and MBC-derived mAbs. We found similar frequencies of heavy chain mutations in antibodies from both cell types (Supp. Fig. 1F), consistent with the larger unpaired dataset.

### Plasmablasts and memory B cells produce highly potent antibodies against SARS-CoV-2

We evaluated the potency of the 216 mAbs in neutralizing authentic SARS-CoV-2 in a high-throughput assay. The majority of mAbs were non-neutralizing, but several were potent neutralizers with IC_50_ values in the ng/mL range (Fig. 1C; Supplementary Table 1). Most of the neutralizing antibodies had low mutation levels (<3%) (Fig. 1D). For antibodies that were originally of the IgA isotype, we compared neutralization of both IgA and IgG forms and found that the IgA generally showed superior neutralization (Supp. Fig. 2A)^27, 28^. Of the 21 antibodies with IC_50_ <1 µg/mL (as IgG), 16 targeted the RBD and 5 bound to the NTD, consistent with previous reports describing the RBD as the primary neutralizing site^11, 12, 19^. All antibodies that did not bind to RBD or NTD did not reach the IC_50_ <1 µg/mL neutralization threshold. Of the 21 most potent mAbs, 16 originated from plasmablasts and 5 from MBCs. The average potency of all neutralizing antibodies from both cell types was similar (Fig. 1E), suggesting that newly differentiated plasmablasts and MBCs can both produce potent antibodies. When we only considered antibodies from donor COV050, plasmablast-derived mAbs tended to be more potent than MBC-derived mAbs, but this was not statistically significant (P = 0.0728) (Supp. Fig. 2B).

**Fig. 2.**
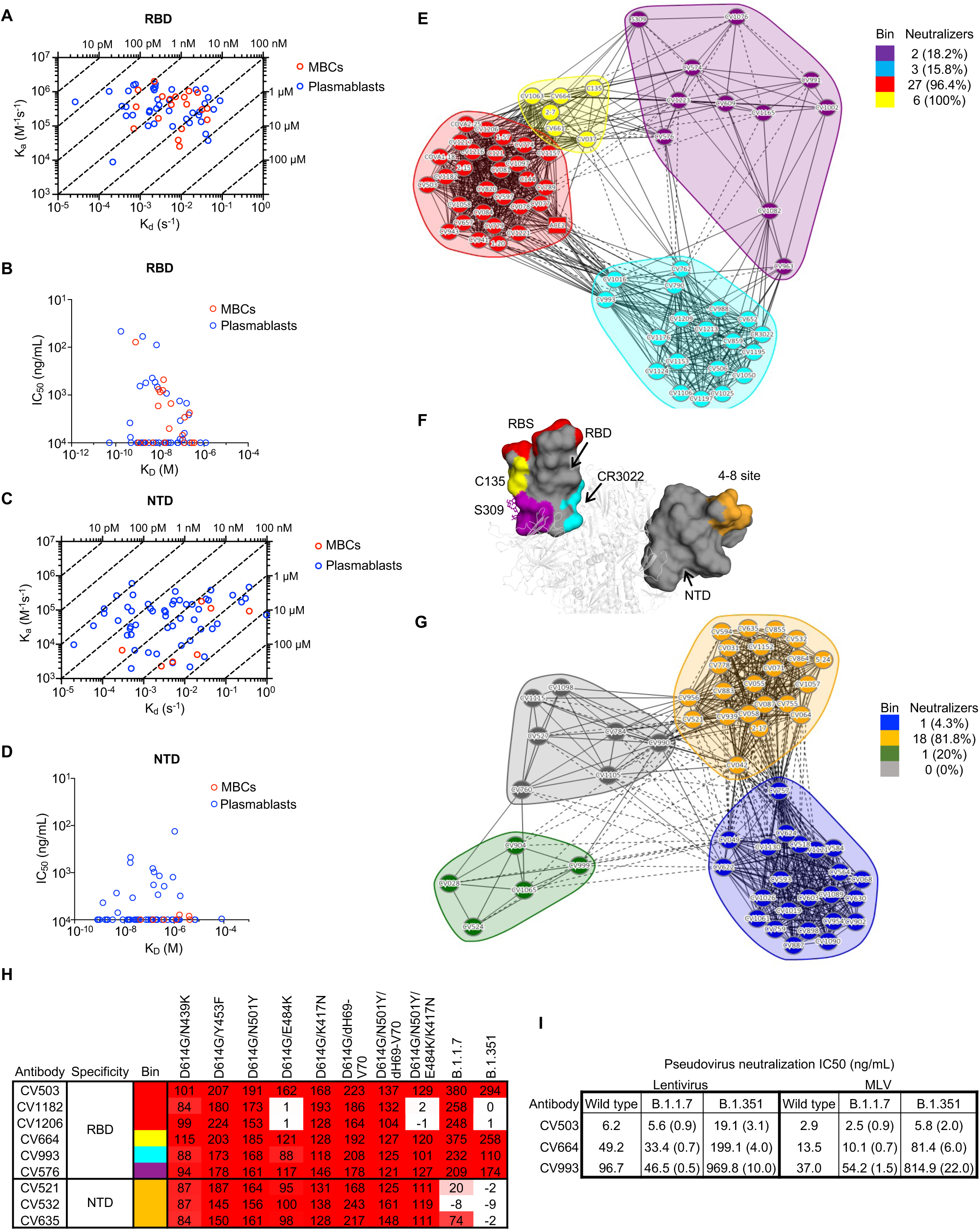
Potent antibodies against SARS-CoV-2 target diverse epitopes on the RBD and NTD and neutralize emerging variants. **A**, Isoaffinity plot of antibodies targeting SARS-CoV-2 RBD (representative of n = 2 experiments). The affinity (KD) values on the top and right of the plot refer to the dotted lines crossing the plot, e.g. any antibody falling on the 10 pM dotted line has a KD value of 10 pM. **B**, Neutralization potency versus affinity of anti-RBD antibodies. **C**, Isoaffinity plot of antibodies targeting SARS-CoV-2 NTD (representative of n = 2 experiments). The affinity (KD) values on the top and right of the plot refer to the dotted lines crossing the plot, e.g. any antibody falling on the 10 pM dotted line has a KD value of 10 pM. **D**, Neutralization potency versus affinity of anti-NTD antibodies. **E**, Epitope binning of anti-RBD antibodies (representative of n = 2 experiments). ACE2 was only used as an analyte (competitor) and not as a ligand, while all other antibodies were tested as both ligands and analytes. Solid lines indicate two-way competition while dotted lines indicate one-way competition. The number and percentage of neutralizing antibodies (IC50 < 10 µg/mL) in each bin are shown. **F**, Epitope bins represented by C135 (yellow bin), S309 (purple bin), ACE2 (red bin), CR3022 (cyan bin), as well as the NTD-specific antibody 4-8 (orange) modeled onto a SARS-CoV- 2 spike protein (white cartoon). Antibody 4-8 was not binned successfully in our experiments but binds to a similar region to 2-17 and 5-24 (ref. 5), which were binned. The epitope sites are color-coded the same as in Fig. 2E and 2G. N-glycans at the N343 glycan site are represented by sticks. **G**, Epitope binning of anti-NTD antibodies (representative of n = 2 experiments). All antibodies were tested as both ligands and analytes. Solid lines indicate two-way competition while dotted lines indicate one- way competition. The number and percentage of neutralizing antibodies (IC50 < 10 µg/mL) in each bin are shown. **H**, Binding of mAb panel to spike protein containing mutations from B.1.1.7 and B.1.351 variants (n = 1 experiment). The numbers show the percentages of mAb binding to mutants relative to D614G (which was normalized to 100). **I**, Neutralization potency of CV503, CV664 and CV993 against B.1.1.7 and B.1.351 variants relative to wild-type (pseudotyped) SARS-CoV-2 (n = 1 experiment). Ratios are shown in brackets, and numbers smaller than 1 indicate an increase in potency while numbers larger than 1 indicate a decrease in potency relative to wild-type.

To determine the relative potency of these mAbs compared to highly potent mAbs described by others^11, 18, 21^, we expressed 10 benchmark IgG1 mAbs: C121, C135, C144 from (ref. 21), COVA1- 18 and COVA2-25 from (ref. 18), and 2-15, 2-7, 1-57, 2-17 and 5-24 from (ref. 11). To enable an accurate comparison of their relative potency, we performed three different neutralization assays: two with authentic SARS-CoV-2 and one with a SARS-CoV-2 pseudotyped virus (Fig. 1F,G and Supp. Fig. 2C,D). Our most potent mAbs, in particular CV503 and CV664, performed comparably to the benchmark mAbs across the different assays. Surprisingly, the NTD-specific antibodies as well as C135^21^, which targets the RBD but does not block ACE2 binding, did not show efficacy in the pseudovirus assay (Fig. 1G). Furthermore, across the three assays, the relative potency of mAbs was reasonably consistent, but absolute IC_50_ values varied greatly (Fig. 1G), highlighting the importance of using standardized assays to compare antibodies from various sources (see CoVIC, covic.lji.org and ACTIV, https://www.nih.gov/research-training/medical-research-initiatives/activ).

### Potent antibodies against SARS-CoV-2 target diverse epitopes on the RBD and NTD

Next, we used high-throughput surface plasmon resonance (SPR) to determine the affinity of the RBD- and NTD-specific mAbs. Overall, RBD-specific mAbs had higher affinity than NTD-specific mAbs (Supp. Fig. 3A). When stratified by antigen specificity and cell type, RBD-specific mAbs derived from plasmablasts and MBCs had similar affinities, with a few antibodies reaching sub-nM affinity (Fig. 2A,B and Supp. Fig. 3B). The top 4 neutralizing mAbs had high affinity (<10 nM) to the RBD, but other high-affinity antibodies, including the antibody with the highest affinity in our panel, were non-neutralizing (Fig. 2B), suggesting that high affinity for the RBD as a soluble protein may be necessary but not sufficient for neutralization in the context of the virus. In contrast, NTD-specific mAbs from plasmablasts generally had lower affinities than those from MBCs (Fig. 2C,D and Supp.. Fig. 3C). Surprisingly, the most potent mAb had relatively low affinity for the NTD, approaching micromolar K_D_ (Fig. 2D). Analysis of antibodies from donor COV050 showed that plasmablast-derived mAbs had higher affinity than MBC-derived mAbs (Supp. Fig. 3D), which is consistent with the neutralization data for this donor, as well as previous work suggesting that higher BCR affinity is associated with differentiation into plasma cells^29–31^.

**Fig. 3.**
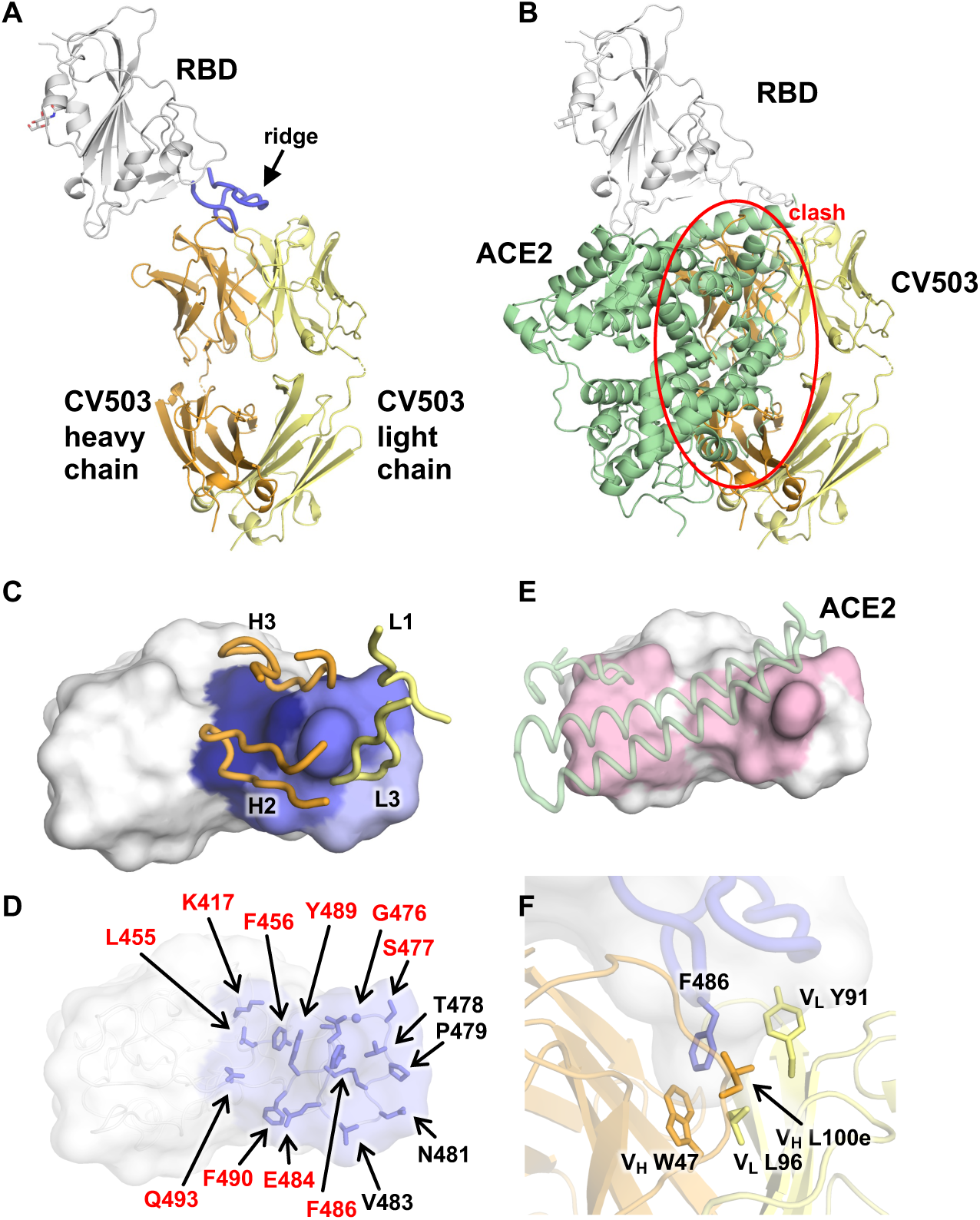
Crystal structure of SARS-CoV-2 RBD in complex with CV503. **A**, CV503 binds to the ridge region of SARS-CoV-2 RBD. The heavy and light chains of CV503 are shown in orange and yellow, respectively. SARS-CoV-2 RBD is in white, where its ridge region (residues 471–491) is shown in blue. **B,** The ACE2/RBD complex structure (PDB ID: 6M0J)^52^ is superimposed on the CV503/RBD complex. The heavy chain of CV503 (orange) would clash with ACE2 (green) if bound to RBD simultaneously (indicated by red circle). **C-D**, Epitope of CV503. Epitope residues contacting the heavy chain are in dark blue and light chain are in light blue, while residues contacting both heavy and light chains are in ocean blue. In **C**, CDR loops that are directly involved in RBD-binding are labeled. In **D**, epitope residues are labeled. Epitope residues that are also involved in ACE2 binding are labeled in red. **E,** ACE2-binding site on the RBD are in light pink. ACE2 is represented as semi-transparent cartoon in pale green. Epitope residues and ACE2-interacting residues are defined as those with a buried surface area (BSA) > 0 Å^2^. **F**, F486 at the ridge region of SARS-CoV-2 RBD (blue) is clamped in a hydrophobic pocket formed by the heavy (orange) and light chains (yellow) of CV503.

We used high-throughput SPR to perform epitope binning of the antibodies and included several antibodies with known binding sites as controls. The RBD-specific antibodies fell broadly into four bins (Fig. 2E,F and Supp. Fig. 4). Most RBD-specific neutralizing antibodies mapped to a bin with ACE2 or a second site containing benchmark antibody C135, which is located on the side of the RBD (Fig. 2F). In contrast, nearly all neutralizing NTD-specific antibodies, including all benchmark NTD-binding antibodies tested^11^, mapped to a single bin (Fig. 2F,G and Supp. Fig. 5), consistent with the recent identification of a single antigenic supersite on the NTD^32, 33^. Interestingly, the three most potent mAbs, CV503, CV664 and CV993, mapped to separate epitope bins and did not compete for binding to RBD. CV503 (red bin) bound to the ACE2 receptor-binding site (RBS), while CV664 (yellow bin) and CV993 (cyan bin) targeted opposing sides of the RBD away from the RBS (Fig. 2F). This finding suggested that some of these antibodies may function against the emerging SARS-CoV-2 variants B.1.1.7 and B.1.351, since antibodies that do not directly target the RBS may be more resistant to variant mutations^8^.

**Fig. 4.**
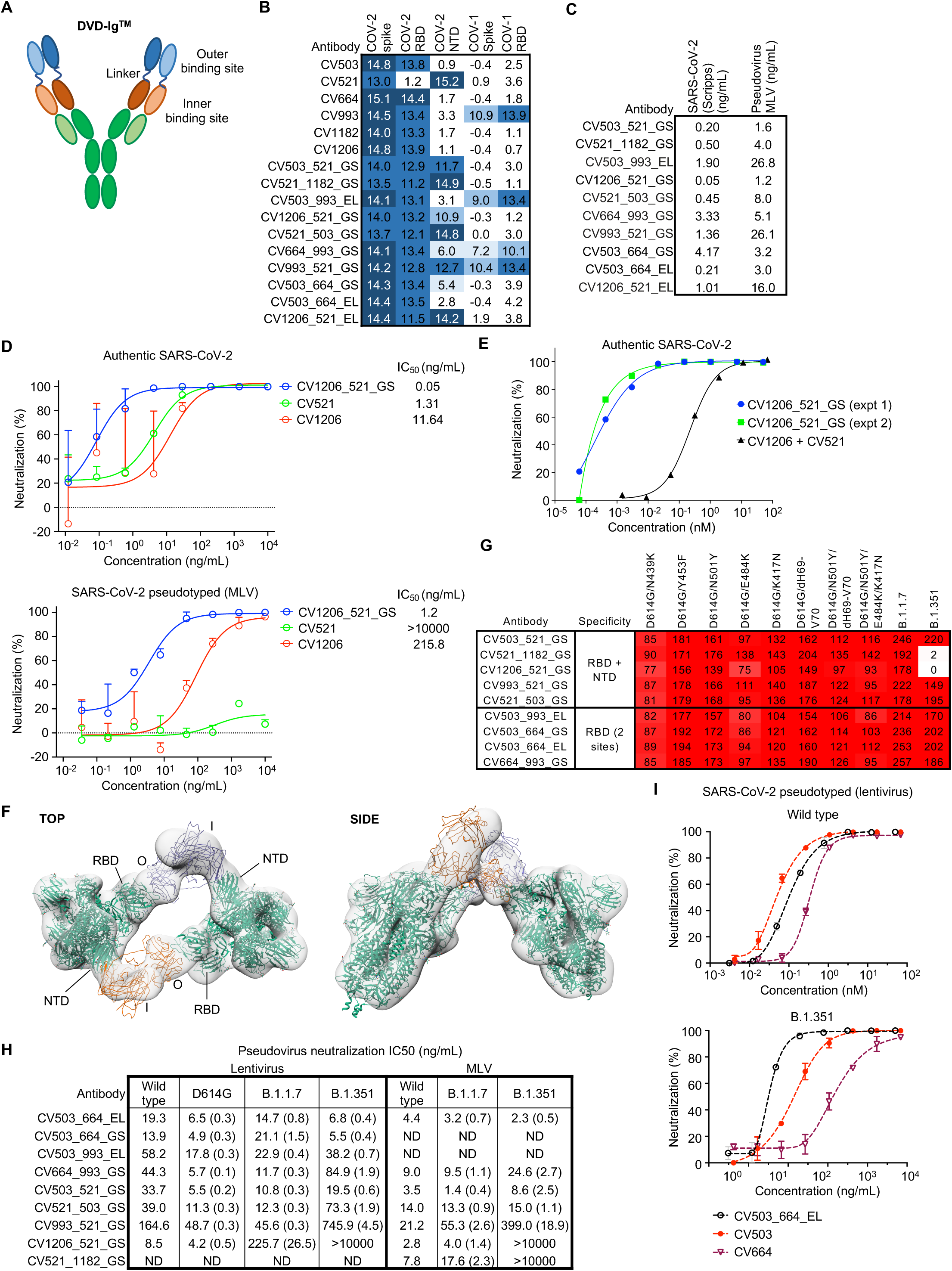
Bispecific antibodies utilize both binding sites to potently neutralize SARS-CoV-2 and are effective against emerging variants. **A**, Scheme of DVD-Ig^TM^. In our bispecific antibody naming system, the first name refers to the antibody used to make the outer binding site and the second refers to the antibody at the inner binding site. GS or EL refers to the type of linker connecting the two antigen-binding sites. See Materials and Methods for details. **B**, Binding of individual and bispecific antibodies to various domains from SARS-CoV-1 and SARS-CoV-2 (representative of n = 2 experiments). Area under the curve (AUC) values are shown after subtraction with the negative control antigen. **C**, Neutralization potency of bispecific antibodies with SARS-CoV-2 authentic and pseudotyped virus (MLV). Values are averaged from two experiments done in duplicate. **D**, Neutralization curves of CV1206_521_GS with SARS-CoV-2 authentic and pseudotyped virus. Curves are from a representative experiment, IC50 values for authentic virus are the average from two experiments and those for the pseudovirus are from an average of two (bispecific) or three (regular antibody) experiments. **E,** Neutralization potency of CV1206_521_GS versus a cocktail of CV1206 and CV521, with concentrations shown in the molar scale to enable a fair comparison. For the antibody combination, the values on the x-axis refers to the concentration of each antibody in the cocktail, e.g. 10 nM refers to 10 nM of CV1206 + 10 nM of CV521. **F**, 3D reconstruction of CV1206_521_GS from nsEM images. Two “one RBD up” models (PDB 6VYB) in green are docked into the reconstruction. Similarly, multiple mock scFv’s in orange and purple were docked to approximate the DVD-Ig molecule. O, outer binding site; I, inner binding site. **G**, Binding of bispecific antibody panel to spike protein containing mutations from B.1.1.7 and B.1.351 variants (n = 1 experiment). The numbers show the percentages of mAb binding to mutants relative to D614G (which was normalized to 100). **H**, Neutralization potency of bispecific antibodies against D614G, B.1.1.7 and B.1.351 variants relative to wild-type (pseudotyped) SARS-CoV-2 (n = 1 experiment). Ratios are shown in brackets: numbers smaller than 1 indicate an increase in potency while numbers larger than 1 indicate a decrease in potency relative to wild-type. ND, not determined. **I,** Potency of CV503_664_EL versus individual component mAbs against wild-type and B.1.351 SARS-CoV-2 pseudotyped virus (lentivirus).

**Fig. 5.**
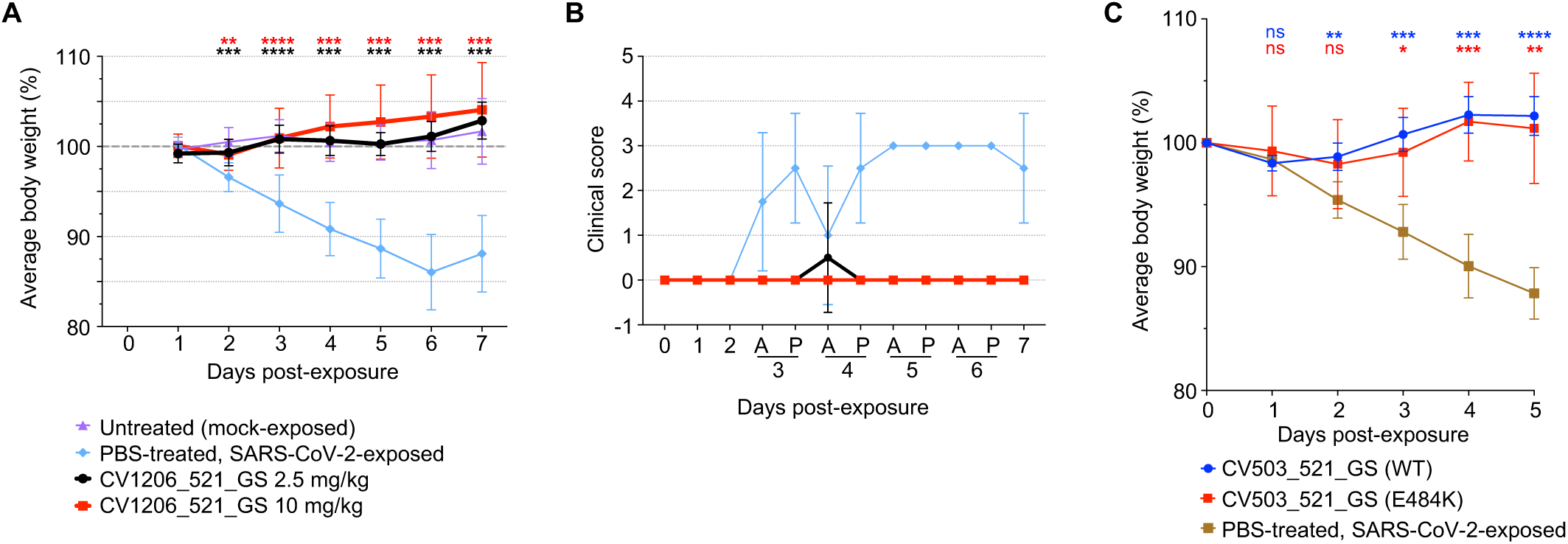
Bispecific antibodies prevent disease mediated by WT or E484K SARS-CoV-2 in an *in vivo* model. **A,** Weight change in hamsters that were administered CV1206_521_GS IP at a dose of 2.5 or 10 mg/kg, 24 h prior to IN virus exposure at 5log10 PFU. Differences between groups that were given the antibody versus PBS were determined using a mixed-effects repeated measures analysis with Dunnett’s multiple comparisons; **P<0.01, ***P<0.001, ****P<0.0001. n = 12 hamsters per group for days 0-3, n = 6 per group for days 4-7. Points represent mean +/- standard deviation (SD). **B,** Blinded clinical score assigned to hamsters throughout the course of disease. A, AM; P, PM. **C,** Weight change in hamsters that were administered bispecific antibodies at 1 mg/hamster, 12 h prior to IN virus exposure at 5log10 PFU. Differences between groups that were given the antibody versus PBS were determined using a mixed-effects repeated measures analysis with Dunnett’s multiple comparisons; *P<0.05, **P<0.01, ***P<0.001, ****P<0.0001. n = 5 hamsters per group. Points represent mean +/- standard deviation (SD).

We tested the ability of all neutralizing antibodies to bind to B.1.1.7 and B.1.351 spike protein (Supp. Fig. 6A). Twenty-eight out of 37 RBD-specific mAbs and 3 out of 20 NTD-specific mAbs retained full (>90%) binding to B.1.1.7, but only 10 RBD-specific mAbs and 2 NTD-specific mAbs retained full binding to B.1.351, consistent with previous findings suggesting that the B.1.351 variant is more successful than B.1.1.7 in evading the neutralizing antibody response^7, 8^.

We narrowed our focus to potent antibodies that targeted distinct epitope bins and screened them against a larger panel of SARS-CoV-2 mutants encoded within B.1.1.7 and B.1.351 to enable identification of specific deleterious mutations (Fig. 2H). CV664 and CV993, along with CV576, which mapped to the 4^th^ RBD bin (purple), retained binding to all B.1.1.7 and B.1.351 spike protein variants. Surprisingly, our most potent antibody CV503 (RBS-specific) also retained binding to all variants tested, while CV1182 and CV1206, which share a bin with this antibody, failed to bind to B.1.351 due to the E484K mutation. Consistent with these findings, CV503, CV664 and CV993 were able to neutralize B.1.1.7 and B.1.351 (Fig. 2I). All three antibodies neutralized the B.1.1.7 variant with no loss in potency relative to wild-type. CV503 neutralized the B.1.351 variant with a slight (2- to 3-fold) loss in potency, while CV664 and CV993 had a 4- to 22-fold reduction in potency against this variant. In contrast, all NTD-specific antibodies failed to bind to the B.1.351 spike, and two out of three also had sharply reduced or abolished binding to B.1.1.7 (Fig. 2H), consistent with previous work suggesting that antibodies targeting the NTD supersite are susceptible to variant mutations^8^.

### Crystal structure of CV503 reveals a binding site that overlaps with ACE2 with limited interactions with key mutant residues

To understand the mechanism that enabled CV503 to retain binding to the SARS-CoV-2 variants, we determined the crystal structure of Fab CV503 in complex with SARS-CoV-2 RBD at 3.4 Å resolution (Fig. 3, Supplementary Table 2). Another SARS-CoV-2 targeting Fab, COVA1-16, that binds a different site on the RBD^18, 34^, was co-crystallized to enhance crystal lattice formation (Supp. Fig. 6B). The X-ray structure confirmed that CV503 binds to the RBS of SARS-CoV-2. The buried surface area (BSA) of SARS-CoV-2 RBD conferred by the heavy and light chains of CV503 are 487 Å^2^ and 290 Å^2^, respectively. CV503 almost exclusively targets the ridge region (residues 471–491) of the RBS, with 80% of the antibody interaction focused on this region (Fig. 3A). In fact, the ridge region of SARS-CoV-2 RBD dominates the epitope area for several highly potent monoclonal antibodies in addition to CV503, e.g. S2E12 where the ridge occupies 80% of the interaction area of the RBD (PDB ID: 7K45)^20^, P17 (79% of area, PDB ID: 7CWO)^35^ and COVOX-253 (74% of area, PDB ID: 7BEN)^36^. These ridge-dominated neutralizing antibodies are encoded by diverse germlines, including VH1-69, VH3-30, and VH1-58 that encode structurally convergent RBD-targeting antibodies^36^. This finding highlights the highly immunogenic ridge region as a highly vulnerable area on the SARS-CoV-2 RBD. CV503 clearly interferes with the binding of ACE2 to the RBD (Fig. 3B-E). Notably, F486 at the tip of the ridge region is inserted into a hydrophobic pocket formed by the heavy (V_H_ W47 and L100e) and light (V_L_ L96 and Y91) chains (Fig. 3F). This interaction is similar to those formed by S2E12^20^ and COVOX-253^36^, which also form hydrophobic pockets with six aromatic amino acids from both heavy and light chains that stack with RBD-F486 (Supp. Fig. 6C-J). Similarly, P17 also stacks with RBD-F486 with two tyrosines^35^. Although both target the same area, S2E12 and COVOX-253 are rotated 50° compared to CV503, whereas P17 is rotated by 85° (Supp. Fig. 6G). CV503 does not form extensive interactions with K417 and E484, which are located on the periphery of the epitope (Fig. 3D), and does not interact with N501, explaining the resistance of this antibody to mutations in the B.1.1.7 and B.1.351 variants.

### Bispecific antibodies potently neutralize SARS-CoV-2

Given that the most potent mAbs bound to non-overlapping sites, we assessed possible synergy between these antibodies. We tested RBD-specific antibodies CV503, CV664, CV993 and CV1182, along with the most potent NTD-binder (and 5^th^ best overall neutralizer) CV521 in an initial screen with pairwise combinations between all non-competing antibodies (Supp. Fig. 7A). Most combinations, including all RBD-NTD pairs, were additive or inhibitory, but two RBD-specific pairs, CV503+CV664 and CV664+CV993, showed evidence of synergy at two different concentrations. However, follow-up tests with a larger range of concentrations gave inconsistent results, making it unclear if these antibodies were truly synergistic (Supp. Fig. 7B,C). We then hypothesized that merging multiple antibody specificities in the same molecule might provide a different effect than simply mixing two antibodies. Thus, we designed and produced bispecific DVD-Ig^TM^ antibodies combining the variable regions of the potent neutralizers with two types of linkers (GS or EL, see Materials and Methods)^37^. This form of a bispecific antibody is relatively easy to express and has a similar structure to standard IgG, except for the addition of an antigen-binding domain on top (outer domain) of the native binding domain (inner domain) through a flexible linker (Fig. 4A). We successfully expressed 10 DVD-Ig^TM^ antibodies, and SDS-PAGE and size exclusion chromatography confirmed that 9 of 10 antibodies contained a single dominant product with the expected molecular weight (Supp. Fig. 8A,B). We found that bispecific antibodies that combined an RBD-specific and NTD-specific antibody retained binding to both domains (Fig. 4B). Moreover, SPR experiments confirmed that bispecific antibodies containing two different RBD-binding sites were able to utilize both sites (Supp. Fig. 8C). For instance, CV503_664_GS, which combined CV503 and CV664, was able to bind RBD previously attached to either component antibody (Supp. Fig. 8C). Negative stain electron microscopy (nsEM) revealed that the 10 bispecific antibodies each cross-linked 2 to 4 spike proteins (Supp. Fig. 9A), and 3 of 10 antibody-spike complexes were amenable to 3D refinement (Supp. Fig. 9B). We tested the panel of bispecific antibodies in authentic SARS-CoV-2 and pseudovirus neutralization assays (Fig. 4C).

Five bispecific antibodies, CV503_521_GS, CV521_1182_GS, CV1206_521_GS, CV521_503_GS and CV503_664_EL, were able to neutralize authentic SARS-CoV-2 with an IC_50_ of <1 ng/mL (Fig. 4C). CV1206_521_GS was the most potent neutralizing antibody in our panel (either bispecific or native IgG), which was surprising given the lower potency of antibody CV1206 compared to other RBD-specific antibodies (Fig. 1G, 4D). Notably, CV1206_521_GS neutralizes SARS-CoV-2 with >100-fold higher potency than a cocktail of its constituent antibodies CV1206 and CV521 (Fig. 4E). We found that CV1206_521_GS uses its inner and outer Fab domains to cross-link NTD and RBD in adjacent spike proteins (Fig. 4F), a mode of action that is unavailable to conventional mAbs, even when used in combination. Taken together, these data suggest that pairing suitable antibodies in a bispecific format can dramatically improve potency by introducing novel mechanisms of action.

### Bispecific antibodies are resistant to mutations in emerging SARS-CoV-2 variants

Another potential advantage of bispecific antibodies is resistance to current and future viral escape mutants, since these antibodies target multiple sites on the spike protein. We tested the bispecific antibodies for binding to variant spike proteins carrying individual and total mutations encoded by B.1.1.7 and B.1.351. Only the bispecific antibodies whose two components both lost binding to the SARS-CoV-2 variants were unable to bind to these variants (Fig. 4G), suggesting that this antibody form is more resistant to spike mutations than regular mAbs. To confirm the binding results, we tested the ability of nine bispecific antibodies to neutralize SARS-CoV-2 D614G, B.1.1.7 and B.1.351 pseudotyped virus in two independent labs. All the bispecific antibodies neutralized D614G with no loss of efficacy (Fig. 4H). All dual-RBD binders and most bispecific antibodies that contained CV521 (which had sharply reduced binding to B.1.1.7 as a mAb) effectively neutralized the B.1.1.7 variant with equal potency compared to wild-type virus (Fig. 4H). Importantly, six out of nine bispecific antibodies tested neutralized B.1.351 similarly (within 3-fold variation) to wild-type SARS-CoV-2, despite the complete loss of binding of one component (CV521) in two of these antibodies and the reduced potency of at least one component (CV503/CV664/CV993) in all the others (Fig. 4H). For instance, a bispecific antibody combining CV503 and CV664 (CV503_664_EL) actually improved slightly in potency against the B.1.351 variant, even though CV503 and CV664 were both less effective against the variant as individual mAbs (Fig. 4I, Supp. Fig. 10A). In addition, an equimolar cocktail of CV503 and CV664 (where x molar of CV503 + x molar of CV664 was compared with x molar of CV503_664_EL) was about 2-fold less effective in neutralizing both wild-type and B.1.351 SARS-CoV-2 (Supp. Fig. 10A).

### Bispecific antibodies are highly effective in an *in vivo* model of SARS-CoV-2 infection

We tested the bispecific antibodies for efficacy against SARS-CoV-2 infection in the well-established Syrian hamster model that resembles features of severe COVID-19 in humans (Fig. 5)^12, 13, 38–41^. In the first experiment, the bispecific antibodies were delivered intraperitonially (IP) at 2.5 mg/kg or 10 mg/kg, followed by intranasal (IN) administration of 10^5^ PFU SARS-CoV-2 24 hours later. Change in body weight and a blinded clinical score were used to assess SARS-CoV- 2-mediated disease. Consistent with previous reports, hamsters injected with PBS as sham treatment had a >10% reduction in body weight through day 6 post-infection, followed by a rebound in weight, while mock-exposed hamsters had no weight change (Fig. 5A)^38, 39, 41^. In contrast, SARS-CoV-2-exposed hamsters that had received 2.5 mg/kg or 10 mg/kg of CV1206_521_GS had no weight loss through the week-long observation period, similar to the uninfected controls (Fig. 5A). No clinical signs (Supp. Fig. 10B) were observed in hamsters in the mock-exposed group or in the virus-exposed groups that received either dose of CV1205_521_GS, with the exception of one hamster in the 2.5 mg/kg group that had a rapid respiratory rate on day 4, but then recovered (Fig. 5B). In contrast, rapid shallow breathing was observed in 7 of 12 hamsters in the sham-treated SARS-CoV-2-exposed group starting on Day 3, while all remaining hamsters in this group developed clinical signs by Day 5 through the end of the study (Fig. 5B). We confirmed the efficacy of CV1206_521_GS in preventing clinical disease caused by wild-type SARS-CoV-2 in the hamster model at an independent laboratory (Supp. Fig. 10C). Next, we tested the *in vivo* efficacy of a potent bispecific antibody that neutralized B.1351, CV503_521_GS, against SARS-CoV-2 carrying the critical E484K variant mutation, which reduces the neutralization potency of many mAbs and convalescent plasma^4, 8^. Importantly, CV503_521_GS was equally effective *in vivo* against wild-type SARS-CoV-2 and the virus carrying this mutation (Fig. 5C), matching our *in vitro* findings (Fig. 4H). We also tested an equimolar cocktail of the mAbs CV503 + CV521, as well as mAbs CV1206 + CV521, in the same model but could not distinguish between these cocktails and their corresponding bispecific antibodies as both were equally protective at the dose tested (Supp. Fig. 10C). Nevertheless, the *in vitro* neutralization results with CV1206_521_GS (Fig. 4E) suggest that certain bispecific antibodies have higher potency than cocktails of the parent mAbs. Altogether, these results suggest that the bispecific antibodies are effective in preventing SARS-CoV-2-mediated disease *in vivo*, including disease caused by SARS-CoV-2 carrying a key variant mutation.

## Discussion

The emergence of new SARS-CoV-2 variants^1–3^ that appear to compromise vaccine efficacy and are resistant to many existing mAbs, including some currently in the clinic^4–9^, provides strong motivation to broaden the search for potent mAbs against the virus. In this study, we found that both plasmablasts and MBCs are capable of producing high-affinity, potent neutralizing antibodies that target diverse regions of the SARS-CoV-2 spike protein. These findings suggest that plasmablasts are a previously untapped source of potent mAbs targeting SARS-CoV-2 that should be interrogated further. We did not unequivocally find synergy among the antibody combinations tested, consistent with the paucity of studies demonstrating true synergy among SARS-CoV-2- specific mAbs^10^. Nevertheless, antibody cocktails that target different regions of the spike protein offer the potential advantage of being more resistant to emerging variants. Indeed, antibody cocktails are the main format currently used in the clinic for treatment of SARS-CoV-2^42, 43^.

We explored the use of DVD-Ig^TM^ bispecific antibodies as a potential alternative to antibody cocktails. Bispecific antibodies have recently been approved by the FDA for treatment of hemophilia A^44^ and acute lymphoblastic leukemia^45^, and are under investigation for other malignancies^46^ and for HIV^47–49^. Bispecific antibodies offer three potential advantages over mAbs or mAb cocktails. First, bispecific antibodies can neutralize SARS-CoV-2 via mechanisms not available to mAbs, as demonstrated by the ability of CV1206_521_GS to cross-link adjacent spike proteins using its dual RBD and NTD specificities, which was associated with superior neutralization compared to a cocktail of the parent mAbs CV1206 and CV521. Second, bispecific antibodies are more resistant to emerging SARS-CoV-2 variants than single mAbs, since they target multiple sites of the spike protein. Indeed, six of nine bispecific antibodies tested retained the ability to neutralize the B.1.351 variant at near wild-type potency, even though all six bispecific antibodies contained at least one antibody component that showed reduced binding or neutralization of B.1.351 as a mAb. Crucially, a potent bispecific antibody was efficacious against SARS-CoV-2 carrying the key variant mutation E484K in the hamster model. Third, from a clinical development standpoint, as single molecules, bispecific antibodies potentially offer practical and cost advantages over mAb cocktails^50^. In the face of rapidly emerging SARS-CoV-2 variants of concern, our findings support the further exploration of bispecific antibodies that strategically combine antibody pairs^51^ as new tools in the fight against COVID-19.

## Methods

### Cohort and biospecimens

New York Blood Center (NYBC) collected plasma and whole blood from donors who had been diagnosed with symptomatic SARS-CoV-2 infection by RT-PCR and were without COVID-19 symptoms for at least two weeks at the time of blood collection, according to FDA guidance. All donors signed the standard NYBC blood donor consent form that indicates blood may be used for research. Whole blood remaining after infectious disease testing (3-7 mL/donor) was deidentified and provided for research purposes. Eligible donors were adults ≥18 years of age. All blood samples were collected during the month of April 2020. Peripheral blood mononuclear cells (PBMCs) were isolated by Ficoll-Paque density gradient centrifugation. Following centrifugation, 1-2 mL of plasma from the top layer was removed, transferred to Cryovial tubes, and stored at -80°C. PBMCs were recovered and washed with Dulbecco’s PBS and resuspended in 1 mL of Cryostor CS-10. PBMCs were kept at -80°C for 24 h, and then stored at -196°C in vapor phase liquid nitrogen. PBMC and plasma samples were transported by courier to the NIH on dry ice (- 78.5°C) where they were stored in liquid nitrogen vapor phase and at -80°C, respectively.

### SARS-CoV-2 and SARS-CoV-1 protein antigens

The SARS-CoV-2 N-terminal domain (NTD) (residues 14–305), receptor-binding domain (RBD) (residues 319–541), and ectodomain of the spike (S) protein (residues 14–1213 with R682G/R683G/R685G/K986P/V987P mutations) (GenBank: QHD43416.1), were cloned into a customized pFastBac vector. The SARS-CoV RBD (residues 306–527) and ectodomain of the spike protein (residues 14–1195, with K968P/V969P mutations (GenBank: ABF65836.1) were similarly cloned. The RBD and NTD constructs were fused with an N-terminal gp67 signal peptide and a C-terminal His_6_ tag. The spike ectodomain constructs were fused with an N-terminal gp67 signal peptide and a C-terminal BirA biotinylation site, thrombin cleavage site, trimerization domain, and His_6_ tag. Recombinant bacmid DNA was generated using the Bac-to-Bac system (Life Technologies). Baculovirus was generated by transfecting purified bacmid DNA into Sf9 cells using FuGENE HD (Promega), and subsequently used to infect suspension cultures of High Five cells (Life Technologies) at an MOI of 5 to 10. Infected High Five cells were incubated at 28 °C with shaking at 110 r.p.m. for 72 h for protein expression. The supernatant was then concentrated using a Centramate cassette (10 kDa MW cutoff for RBD and NTD, 30 kDa MW cutoff for spike proteins, Pall Corporation). The proteins were purified by Ni-NTA, followed by size exclusion chromatography.

### Plasma binding to SARS-CoV-2 and other coronavirus antigens

Streptavidin-coated beads with different intensities of PE-channel fluorescence (Spherotech SVFA-2558-6K and SVFB-2558-6K) were incubated with the following biotinylated antigens: 10 µg/mL MERS spike, NL63 spike, 229E spike, HKU1 spike, OC43 spike (all from Sino Biological), SARS-CoV-2 spike, SARS-CoV-2 RBD, SARS-CoV-2 NTD, SARS-CoV-1 spike, SARS-CoV-1 RBD or 10 µg/mL CD4 as a control. Excess streptavidin sites were blocked with 10 µg/mL of CD4 and the beads were washed and mixed. The beads were stained with 1/50, 1/250 or 1/1250 plasma for 30 min at room temperature, washed, and stained with 2.5 µg/mL goat anti-human IgG Alexa Fluor 647 (Jackson Immunoresearch 109-606-170). The samples were read with the iQue Screener Plus (Intellicyt) high-throughput flow cytometer and FACS data were analysed with Flowjo. Data from titrations were analysed by calculating area under the curve (AUC) for the titration and subtracting the AUC of the negative control antigen.

### Fluorescence Reduction Neutralization Assay (FRNA)

Assays for determining neutralizing titers with authentic SARS-CoV-2 (2019-nCoV/USA-WA1- A12/2020 from the US Centers for Disease Control and Prevention, Atlanta, GA, USA) were performed at the NIH-NIAID Integrated Research Facility at Fort Detrick, MD, USA using a fluorescence reduction neutralization assay (FRNA), as previously described^53^. This assay was performed by incubating a fixed volume of virus [0.5 multiplicity of infection (MOI)] with the antibodies for 1 h at 37 °C prior to adding to Vero E6 cells (BEI, Manassas, VA, USA, NR-596) plated in 96-well plates. Following addition to Vero E6 cells, the virus was allowed to infect the cells and propagate for 24 hours at 37 °C/5% CO_2_, at which time the cells were fixed with neutral buffered formalin. Following fixation, the cells were permeabilized with Triton X-100 and probed with a SARS-CoV/SARS-CoV-2 nucleoprotein-specific rabbit primary antibody (Sino Biological, Wayne, PA, USA, 40143-R001) followed by an Alexa Fluor 647-conjugated secondary antibody (Life Technologies, San Diego, CA, USA, A21245). Cells were counterstained with Hoechst nuclear stain (Life Technologies H3570). Cells in four fields per well were counted with each field containing at least 1000 cells. Each dilution step of test sample(s) was run in quadruplicate. Plates were read and quantified using an Operetta high content imaging system (PerkinElmer, Waltham, MA). Antibodies were screened using a 2-fold serial 12-step dilution. The lower limit of detection was either 1:20 or 1:40 depending upon the dilution series. Assays were controlled using a spike protein specific antibody as positive control in addition to virus-only and uninfected cell controls. Percent neutralization was calculated as 100 – [(percent infected cells in well of interest / average percent infected cells from 6 virus-only wells in matched plate) × 100]. The average percent neutralization of quadruplicate samples was determined per antibody dilution point. Plasma and mAb IC_50_ values were estimated using an automated curve fitting script which fit 4-parameter logistic model nonlinear regressions under 22 different combinations of model constraints and parameter starting values, or if those models failed to converge, a quadratic regression model was used. All 22 combinations of the logistic nonlinear regression were attempted, and if any succeeded, the best fitting model was selected using its R-squared value. Quadratic linear regression was only used if all 22 logistic nonlinear regressions failed. Regardless of the model fit, IC_50_ was estimated using interpolation to find the concentration or dilution that generated 50% neutralization.

### Authentic SARS-CoV-2 neutralization assay (Scripps)

Vero E6 cells were seeded in 96-well half-well plates at approximately 8000 cells/well in a total volume of 50 µl complete DMEM medium (DMEM, supplemented with 10% heat-inactivated serum, 1% GlutaMAX, 1% P/S) the day prior to the in-cell enzyme-linked immunosorbent assay (ELISA). The virus (500 plaque forming units/well) and antibodies were mixed, incubated for 30 minutes, and added to the cells. The transduced cells were incubated at 37°C for 24 hours. Each treatment was tested in duplicate. The medium was removed and disposed of appropriately. Cells were fixed by immersing the plate into 4% formaldehyde for 1 hour before washing 3 times with phosphate buffered saline (PBS). The plate was then either stored at 4℃ or gently shaken for 30 minutes with 100 µL/well of permeabilization buffer (consisting of PBS with 1% Triton-X). All solutions were removed, then 100 µl of 3% bovine serum albumin (BSA) was added, followed by room temperature (RT) incubation for 2 hours.

Primary antibodies against the spike protein were generated from a high-throughput screen of samples from a convalescent, coronavirus disease 2019 cohort (CC)^12^. A mix of primary antibodies consisting of CC6.29, CC6.33, L25-dP06E11, CC12.23, CC12.25, in an equimolar ratio, were used next. The primary antibody mixture was diluted in PBS/1% BSA to a final concentration of 2 µg/ml. The blocking solution was removed and washed thoroughly with wash buffer (PBS with 0.1% Tween-20). The primary antibody mixture, 50 µl/well, was incubated with the cells for 2 hours at RT. The plates were washed 3 times with wash buffer.

Secondary antibody (Jackson ImmunoResearch, Peroxidase AffiniPure Goat Anti-Human IgG (H+L), 109-035-088) diluted to 0.5 mg/ml in PBS/1% BSA was added at 50 µL/well and incubated for 2 hours at RT. The plates were washed 6 times with wash buffer. HRP substrate (Roche, 11582950001) was freshly prepared as follows: Solution A was added to Solution B in a 100:1 ratio and stirred for 15 minutes at RT. The substrate was added at 50 µL/well and chemiluminescence was measured in a microplate luminescence reader (BioTek, Synergy 2).

The following method was used to calculate the percentage neutralization of SARS-CoV-2. First, we plotted a standard curve of serially diluted virus (3000, 1000, 333, 111, 37, 12, 4, 1 PFU) versus RLU using four-parameter logistic regression (GraphPad Prism ver. 8) below:

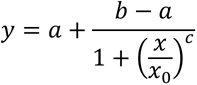

(y: RLU, x: PFU, a,b,c and x_0_ are parameters fitted by standard curve)

To convert sample RLU into PFU, use the equation below: (if y < a then x = 0)

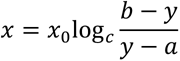

Percentage neutralization was calculated by the following equation:

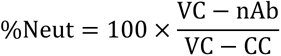

VC = Average of vehicle-treated control; CC = Average of cell only control, nAb, neutralizing antibody. PFU value was used for each variable indicated.

To compute neutralization IC_50_, logistic regression (sigmoidal) curves were fit using GraphPad Prism. Means and standard deviations are displayed in the curve fit graphs and were also calculated using GraphPad Prism.

### MLV pseudovirus neutralization assay

Pseudovirus (PSV) preparation and assay were performed as previously described with minor modifications^12^. Pseudovirions were generated by co-transfection of HEK293T cells with plasmids encoding MLV-gag/pol, MLV-CMV-Luciferase, and SARS-CoV-2 spike (GenBank: MN908947) with an 18 aa truncation at the C-terminus. Supernatants containing pseudotyped virus were collected 48 hr after transfection and frozen at -80°C for long-term storage. PSV neutralizing assay was carried out as follows. 25 µl of mAbs serially diluted in DMEM with 10% heat-inactivated FBS, 1% Q-max, and 1% P/S were incubated with 25 µl of SARS-CoV-2 PSV at 37°C for 1 h in 96-well half-well plates (Corning, 3688). After the incubation, 10,000 Hela-hACE2 cells generated by lentivirus transduction of wild-type Hela cells and enriched by fluorescence-activated cell sorting (FACS) using biotinylated SARS-CoV-2 RBD conjugated with streptavidin-Alexa Fluor 647 (ThermoFisher Scientific, S32357) were added to the mixture with 20 µg/ml Dextran (Sigma, 93556-1G) for enhanced infectivity. At 48 hr post incubation, the supernatant was aspirated, and HeLa-hACE2 cells were then lysed in luciferase lysis buffer (25 mM Gly-Gly pH 7.8, 15 mM MgSO4, 4 mM EGTA, 1% Triton X-100). Bright-Glo (Promega, PR-E2620) was added to the mixture following the manufacturer’s instruction, and luciferase expression was read using a luminometer. Patient samples were tested in duplicate, and assays were repeated at least twice for confirmation. Neutralization ID_50_ titers or IC_50_ values were calculated using “One-Site Fit LogIC50” regression in GraphPad Prism 9.

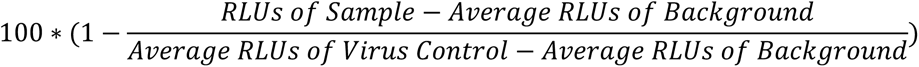

### SARS-CoV-2 variant plasmid generation

The B.1.1.7 plasmid was generated from the pCDNA3.3-SARS2-spike-WT(D18) plasmid with the mutations and primers shown below. Each fragment containing two mutations on the flanking side was PCR-amplified, purified by gel electrophoresis, and then PCR-amplified into one long piece and ligated into the pCDNA3.3 backbone digested with BamHI and XhoI.

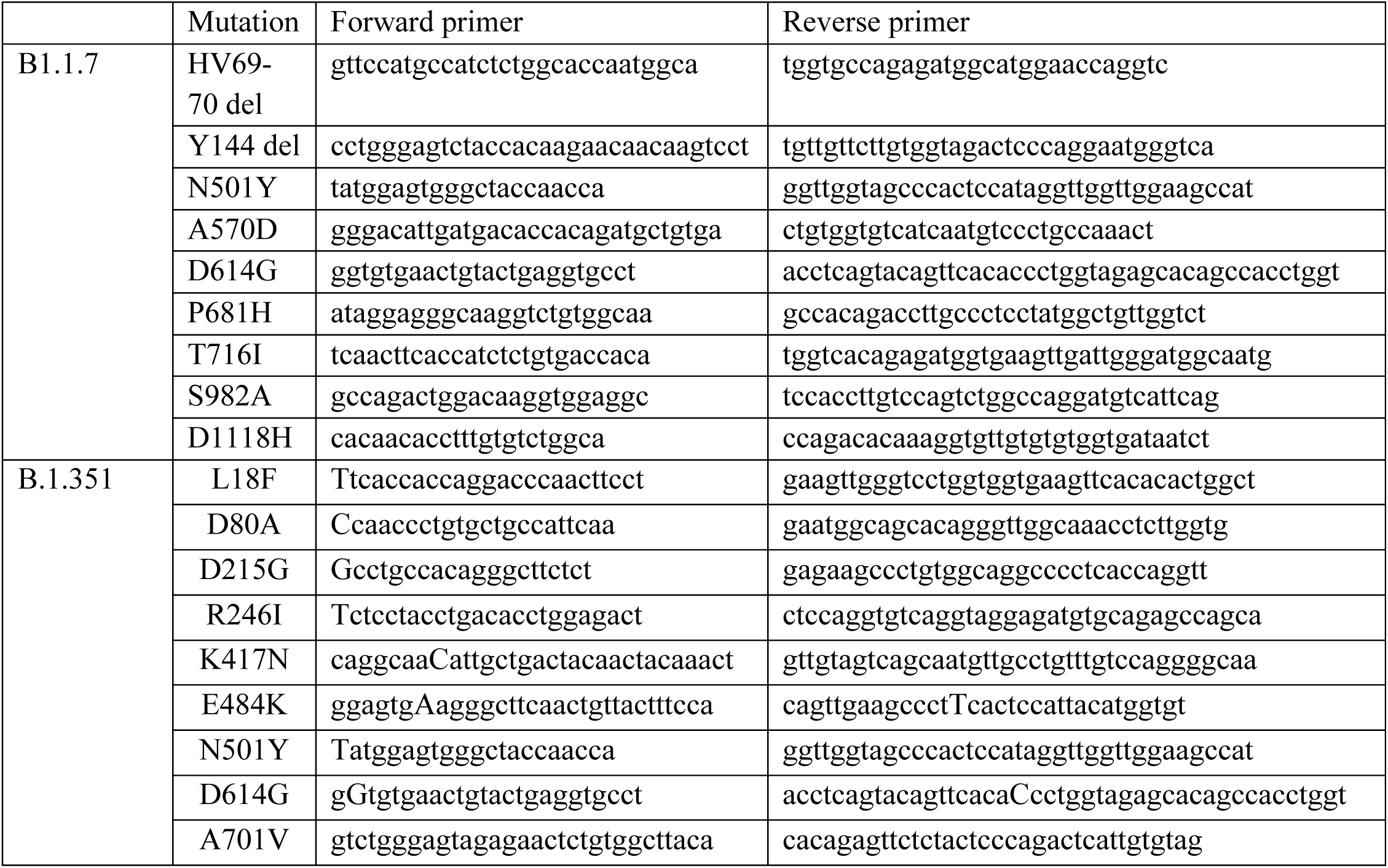

### Lentivirus neutralization assay

S-containing lentiviral pseudovirions were produced by co-transfection of packaging plasmid pCMVdR8.2, transducing plasmid pHR’ CMV-Luc, S plasmid from SARS-CoV-2 (Wuhan-1, D614G, B.1.1.7, B.1.351) with TMPRSS2 into 293T cells using Fugene 6 transfection reagent (Promega, Madison, WI)^54–56^. B.1.1.7 virus contained the following spike mutations: del-H69–V70, del-Y144, N501Y, A570D, D614G, P681H, T716I, S982A, D1118H. B.1.351 virus contained the following spike mutations: L18F, D80A, D215G, del- L242_244, R246I, K417N, E484K, N501Y, D614G, A701V. 293T-ACE2 cells, provided by Dr.

Michael Farzan, were plated into 96-well white/black Isoplates (PerkinElmer, Waltham, MA) at 5,000 cells per well the day before transdution of SARS-CoV-2. Serial dilutions of mAbs were mixed with titrated pseudovirus, incubated for 45 minutes at 37°C and added to 293T-ACE2 cells in triplicate. Following 2 h of incubation, wells were replenished with 100 ml of fresh media. Cells were lysed 72 h later, and luciferase activity was measured with Microbeta (Perking Elmer). Percent neutralization and neutralization IC50s were calculated using GraphPad Prism 8.0.2.

### Antibody binding to cell surface expressed full length SARS-CoV-2 spike proteins

HEK293T cells were transiently transfected with plasmids encoding full length SARS-CoV-2 spike variants using lipofectamine 3000 (L3000-001, ThermoFisher) following manufacturer’s protocol. After 40 hours, the cells were harvested and incubated with bispecific and monoclonal antibodies (1 ug/ml) for 30 minutes. After incubation with the antibodies, the cells were washed and incubated with an allophycocyanin conjugated anti-human IgG (709-136-149, Jackson Immunoresearch Laboratories) for another 30 minutes. The cells were then washed and fixed with 1% paraformaldehyde (15712-S, Electron Microscopy Sciences). The samples were then acquired in a BD LSRFortessa X-50 flow cytometer (BD biosciences) and analyzed using Flowjo (BD biosciences).

### Optofluidics-based identification of SARS-CoV-2-specific mAbs

Cryopreserved PBMCs were thawed and stained with the following panel: LIVE/DEAD Fixable Aqua (ThermoFisher Scientific L34966) or DAPI (BD564907), CD14-BV510 (BioLegend 301842), CD3-BV510 (BioLegend 317332), CD56-BV510 (BioLegend 318340), CD19-ECD (Beckman Coulter IM2708U), IgA-Alexa Fluor 647 (Jackson Immunoresearch 109-606-011), IgD-PE-Cy7 (BD 561314), and IgM-PerCP-Cy5.5 (BD561285), CD27-Alexa Fluor 488 (BioLegend 393204) and CD38-APC-Cy7 (BioLegend 303534). The cells were sorted using the BD FACSAria IIIu in a BSL3 facility and gated on live CD19^+^CD14^-^CD3^-^CD56^-^CD27^++^CD38^++^ (plasmablasts) or live CD19^+^CD14^-^CD3^-^CD56^-^CD27^+^IgM^-^IgD^-^IgA^+^/IgA^-^ (memory B cells). Plasmablasts were sorted into Plasma Cell Survival Medium (Berkeley Lights) and screened in the Beacon device (Berkeley Lights) using standard workflows in the Cell Analysis Suite program. Briefly, sorted plasmablasts were loaded onto an OptoSelect 11k chip and transported as single cells using OEP light cage technology into nanoliter-volume pens. The cells were screened in a 30 min time-course assay for secretion of antibodies that bound to 7 µm streptavidin beads (Spherotech) coated with 10 µg/mL SARS-CoV-2 spike or RBD. Antibody binding to the beads was detected with 2.5 µg/mL goat anti-human IgG-Alexa Fluor 647 (Jackson Immunoresearch 109-606-170), goat anti-human IgA-Cy3 (Jackson Immunoresearch 109-166-011) and goat anti-human IgM-Alexa Fluor 488 (Jackson Immunoresearch 109-546-129), which were added along with the beads during the assay. B cells of interest were selected for RNA extraction and production of cDNA in two ways. First, under the Cell Unload protocol, B cells of interest were exported individually using OEP light cages into 96-well plates for single-cell reverse transcription PCR and total cDNA amplification with the Opto Plasma B Discovery cDNA Synthesis Kit (Berkeley Lights). Alternatively, the BCR Unload protocol was used, where RNA capture beads were imported into the nanopens containing B cells of interest, followed by lysis of the cells, reverse transcription *in situ* to generate cDNA on the beads. The RNA capture beads were exported from the Beacon for total cDNA amplification off-chip. After cDNA amplification from either approach, gene-specific PCR was performed as previously described^57^. Memory B cells were plated into 96-well plates at 2500 cells/well and cultured in a proprietary cytokine cocktail (Berkeley Lights) for 6 days, followed by screening of supernatants for binding to beads coated with 10 µg/mL SARS-CoV-2 spike or RBD using the iQue Screener. B cells from wells of interest were loaded onto the Beacon for single-cell screening, using the same workflows described above.

### mAb sequence analysis and production

Antibody heavy and light chains were PCR-amplified and sequenced as previously described^57, 58^. Sequence analysis, including the determination of the *VH* and *VL* genes and percentage of somatic mutations, was performed using the International Immunogenetics Information System (IMGT) database^59^. Antibody V_H_ or V_L_ sequences were cloned into plasmids containing an IgG1 or relevant light chain backbone (Genscript) and used to transfect Expi293 cells (ThermoFisher Scientific). Recombinant IgG was purified using HiTrap Protein A columns (GE Healthcare Life Sciences). For IgA1 or IgA2 antibodies, V_H_ sequences were also cloned into plasmids containing the IgA1 or IgA2 constant region (Genscript). Recombinant IgA was expressed without a J chain (to express only monomeric IgA) and purified using columns containing the CaptureSelect IgA Affinity Matrix (ThermoFisher Scientific). To produce antibody Fabs, heavy chain plasmids encoding only the V_H_ and C_H_1 were synthesized and used to transfect Expi293 cells along with light chain plasmids. Fab purification (including for the CV503 crystal structure) was performed with the CaptureSelect^TM^ KappaXP Affinity Matrix or CaptureSelect™ LC-lambda (Hu) Affinity Matrix (ThermoFisher Scientific).

### Expression and purification of COVA1-16 Fab

The heavy and light chains of COVA1-16 were cloned into phCMV3. The plasmids were transiently co-transfected into ExpiCHO cells at a ratio of 2:1 (HC:LC) using ExpiFectamine™ CHO Reagent (Thermo Fisher Scientific) according to the manufacturer’s instructions. The supernatant was collected at 10 days post-transfection. The Fabs were purified with a CaptureSelect™ CH1-XL Affinity Matrix (Thermo Fisher Scientific) followed by size exclusion chromatography on a HiLoad Superdex 200 pg column (GE Healthcare), and buffer exchanged into 20 mM Tris-HCl pH 7.4 and 150 mM NaCl.

### mAb binding to SARS-CoV-2 and other coronavirus antigens

For screening of binding to different coronavirus antigens, streptavidin beads were coated as described above with the following biotinylated antigens: 10 µg/mL MERS spike (Sino Biological, 40069-V08B, NL63 spike (Sino Biological, 40604-V08B), 229E spike (Sino Biological, 40605- V08B), HKU1 spike (Sino Biological, 40606-V08B), OC43 spike (Sino Biological, 40607-V08B), SARS-CoV-2 NTD, SARS-CoV-1 spike, SARS-CoV-1 RBD (see above for details of production) or 10 µg/mL CD4 as a control. The beads were stained with 10 µg/mL of each antibody for 30 min at room temperature, washed, and stained with 2.5 µg/mL goat anti-human IgG Alexa Fluor 647 (Jackson Immunoresearch 109-606-170). For screening of binding to SARS-CoV-2 spike and RBD, streptavidin beads were coated with 10 µg/mL of SARS-CoV-2 spike, RBD, or CD4^60^ as a negative control. The beads were incubated with twelve dilutions of the mAbs for 30 min at room temperature, washed, and stained with 2.5 µg/mL goat anti-human IgG Alexa Fluor 647. All samples were read with the iQue Screener Plus (Intellicyt) high-throughput flow cytometer and FACS data were analyzed with Flowjo. Data for binding to coronavirus antigens were analyzed by subtracting median fluorescence intensity (MFI) values of the negative control antigen from the MFI values of each antibody. Data for binding to SARS-CoV-2 spike and RBD were analyzed by calculating area under the curve (AUC) for each antibody without subtraction of negative control antigen.

### Antibody kinetics

All kinetics experiments were performed with the Carterra LSA. A HC30M chip (Carterra) was primed with filtered and degassed HBSTE. The chip was activated with a mixture of 400 mM EDC and 100 mM NHS (ThermoFisher Scientific), followed by coupling with 50 µg/mL goat anti-human IgG Fc (Jackson ImmunoResearch, 109-005-098) in 10 mM sodium acetate pH 5.0, and blocking with 1M ethanolamine, pH 8.5. Next, 1 or 5 µg/mL concentration of each antibody in HBSTE was printed onto a discrete spot on the chip. A series of 7 increasing concentrations of monomeric SARS-CoV-2 RBD (up to 500 nM) or NTD (up to 1.5 µM) was added to the antibody spots sequentially; association was performed for 5 minutes and dissociation for 15 minutes to obtain kinetics data. The results were analyzed as non-regenerative kinetics data using the Kinetics Software (Carterra) to obtain association rate (K_a_), dissociation rate (Kd) and dissociation constant (K_D_) values.

### Epitope binning

Epitope binning experiments were performed with the Carterra LSA. A HC200M chip (Carterra) was primed with filtered and degassed HBSTE + 0.05% BSA. The chip was activated as described above, followed by direct coupling with 10 µg/mL of mAbs of interest in pH 4.5 acetate buffer to discrete spots on the chip and blocking with 1M ethanolamine, pH 8.5. Monomeric 50 nM SARS-CoV-2 RBD or 500 nM SARS-CoV-2 NTD was added to the antibody spots, followed by addition of 10 µg/mL of the sandwiching mAb or protein. Regeneration after each sandwiching antibody was performed with 10 mM glycine pH 2.0. Binning data were analyzed using the Epitope Software (Carterra).

### Antibody synergy experiments

Five antibodies, CV503, CV521, CV664, CV993 and CV1182, were tested for synergy in neutralizing authentic SARS-CoV-2 (FRNA assay). In the initial screens, each antibody was tested at twice the estimated neutralization IC_50_ value (Dose 1) and at the estimated IC_50_ value (Dose 2) in various combinations. Two combinations were explored further: CV503 and CV664, as well as CV664 and CV993. For each pair, each antibody was tested at 5 different concentrations in a checkerboard combination arrangement, along with single antibody controls. Three pairs: CV503 and CV664, CV664 and CV993, and CV1206 and CV521, were also further tested for synergy in the authentic SARS-CoV-2 (Scripps) or pseudovirus assays.

### Design of DVD-Ig^TM^ bispecific antibodies

The mAbs CV503, CV521, CV664, CV993, CV1182 and CV1206 were used to create bispecific DVD-Ig^TM^ antibodies. The antibodies were created as previously described^37^ by synthesizing plasmids that had the heavy chains or the light chains of two antibodies in tandem. For antibodies ending in _GS (Ab1_Ab2_GS), both heavy chains and both light chains were connected by a GGGGSGGGGSGGGG linker, with the first antibody making up the outer variable domains and the second antibody making up the internal domains, closer to the Fc. For antibodies ending in _EL (Ab1_Ab2_EL), the linkers were based on elbow regions between the variable region and constant region: for heavy chains, the sequence ASTKGPSVFPLAP was used. For light chains, the linker sequence QPKAAPSVTLFPP was used when the outer variable domain was of the lambda isotype and the sequence TVAAPSVFIFPP was used when the outer variable domain was a kappa isotype.

### SDS-PAGE

For each antibody, 2.5-5 µg was loaded onto a NuPAGE 4-12% gel (ThermoFisher) and run at 100V for 2-3 h. The gel was stained with PageBlue^TM^ Staining Solution (ThermoFisher) and destained with multiple rinses of water. Imaging was done using the LI-COR Odyssey Fc Imaging System, with emission set at 700 nm.

### Size-exclusion chromatography

To test bispecific antibody purity, each of the antibodies was run on a Superdex 200 column (Increase 10/300 GL, Cytiva) with the AKTA pure chromatography system. The column was pre-equilibrated with PBS. 30 µl of the antibody sample (at 0.5 mg/ml) was applied onto the column and the column was eluted with PBS at 0.75 mL/min. Elution profiles were recorded with Unicorn software (Cytiva).

### Structure modeling

Epitope bins represented by C135 (PDB ID: 7K8Z), S309 (PDB ID: 6WPS), ACE2 (PDB ID: 6M0J), CR3022 (PDB ID: 6W41), as well as NTD-specific antibody 4-8, were modeled onto a SARS-CoV-2 spike protein (PDB ID: 7C2L). Residues with a buried surface area (BSA) > 0 Å^2^, as calculated by the PISA program^61^, were used for defining the epitopes. The epitope of antibody 4-8 refers to an approximate area according to (ref. 11) as coordinates of the complex structure are not publicly available.

### Crystallization and structural determination

Expression and purification of SARS-CoV-2 RBD used for crystallization is similar to the method described in the “SARS-CoV-2 and SARS-CoV-1 protein antigens” section, except that a truncated version of the SARS-CoV-2 RBD (residues 333–529) was used. A complex of CV503 with RBD and COVA1-16 was formed by mixing each of the protein components in an equimolar ratio and incubating overnight at 4°C. The protein complex was adjusted to 14 mg/ml and screened for crystallization using the 384 conditions of the JCSG Core Suite (Qiagen) on our robotic CrystalMation system (Rigaku) at Scripps Research. Crystallization trials were set-up by the vapor diffusion method in sitting drops containing 0.1 μl of protein and 0.1 μl of reservoir solution. Optimized crystals were then grown in 0.1 M sodium citrate pH 4.2, 1 M lithium chloride, and 9% (w/v) polyethylene glycol 6000. crystals appeared on day 7, harvested on day 10 by soaking in reservoir solution supplemented with 15% (v/v) ethylene glycol, and then flash cooled and stored in liquid nitrogen until data collection. Diffraction data were collected at cryogenic temperature (100 K) at Stanford Synchrotron Radiation Lightsource (SSRL) on Scripps/Stanford beamline 12- 1 with a beam wavelength of 0.97946 Å, and processed with HKL2000^62^. Structures were solved by molecular replacement using PHASER^63^ with PDB 7JMW for RBD and COVA1-16^34^, whereas the model of CV503 was generated by Repertoire Builder (https://sysimm.ifrec.osakau.ac.jp/rep_builder/)^64^. Iterative model building and refinement were carried out in COOT^65^ and PHENIX^66^, respectively.

### Single-particle negative stain electron microscopy

The bispecific antibodies were incubated with SARS-2 CoV 6P Mut7 at equal molar ratios for 30 minutes at RT. The complexes were diluted to 0.03 mg/ml in 1X TBS pH 7.4 and applied to plasma-cleaned (Argon/Oxygen mix) copper mesh grids. Uranyl formate at 2% was applied to the grid for 55 seconds and then blotted off. Datasets were collected with a FEI Tecnai Spirit (120KeV, 56,000x magnification) paired with a FEI Eagle (4k x 4k) CCD camera. Data collection details include a defocus value -1.5 μm, a pixel size of 2.06 Å per pixel, and a dose of 25 e^-^/Å^2^. Data collection automation was achieved with the Leginon^67^ software and resulting images were stored in the Appion^68^ database. Complexed single particles were picked using DogPicker^69^ and stacked with a box size of 300 pixels. RELION 3.0^70^ was used for 2D and 3D classifications and final refinements. UCSF Chimera^71^ enabled map segmentation and model docking.

### Hamster study (NIH)

The SARS-CoV virus used for these studies was the 2019-nCoV/USA-WA1-A12/2020 isolate of SARS-CoV-2 provided by the US Centers for Disease Control and Prevention (CDC; Atlanta, GA, USA) that was expanded in Vero E6 cells. Virus sequence was consistent with the published sequence for this isolate. Forty-eight golden Syrian hamsters (*Mesocricetus auratus*, about 6 weeks old, individually housed) sourced from Envigo, Haslett, MI were randomly assigned to four groups of 12 animals each with equal numbers of males and females and a randomized treatment order.

The study was blinded to the staff who handled the animals. Due to safety requirements of the biocontainment laboratory, hamsters were chemically restrained with 4–5% isoflurane for all manipulations. A single treatment of each dose (2.5 and 10 mg/kg) of the CV1206_521_GS antibody or PBS was administered to the respective group by intraperitoneal (IP) inoculation 24 hours prior to intranasal (IN) SARS-CoV-2 (or mock-virus) exposure. Groups of hamsters were then exposed intranasally to SARS-CoV-2 diluted in 100μl (total volume split equally between nostrils) of Dulbecco’s Modified Eagle Medium [DMEM] containing 2% heat-inactivated-FBS to the desired concentration at the 5 log_10_ PFU, or mock inoculum (diluent). The inocula were back- titered in real time to determine the dose received. The actual dose was 5.13 log_10_ PFU as quantified by plaque assay. Following exposure, all hamsters were weighed and monitored for up to 7 days. Animals were observed at least once daily by staff with experience in assessing distress in hamsters and assigned a clinical score.

### Hamster study (Scripps)

As previously described^12^, 8-week old Syrian hamsters were given an IP antibody injection 12 hours pre-infection. Hamsters were infected through intranasal installation of 10^5^ total PFU per animal of SARS-CoV-2 (USA-WA1/2020) or recombinant SARS-CoV-2 E484K in 100uL of DMEM. Hamsters were then weighed for the duration of the study. At day-5 post-infection, animals were sacrificed and lungs were harvested for plaque live virus analysis and histology. The research protocol was approved and performed in accordance with Scripps Research IACUC Protocol #20-0003.

### Statistical analysis

Associations between plasma binding to SARS-CoV-2 spike, RBD, NTD, and plasma neutralization, as well as the correlation between plasma and mAb neutralization of the different donors, were evaluated using a two-tailed Spearman’s rank correlation. Comparisons of characteristics between plasmablasts and MBCs were performed with the Mann Whitney U-test. Synergy was assessed using Loewe’s Additivity through the web application SynergyFinder v2.0 (https://synergyfinder.fimm.fi/)72.

## Acknowledgements

We thank the blood donors at the New York Blood Center; Gavin Wright and Nicole Muller- Sienerth (Wright lab, Wellcome Trust Sanger Institute) for providing recombinant CD4; Kathy Zhao, Aurora Fabry-Woods and Vincent Pai (Berkeley Lights), as well as Andrea Murphy, Dinuka Abeydeera and Ira Seferovich (Carterra) for scientific discussion. We thank Louis Huzella, Ricky Adams, Janie Liang, Saurabh Dixit, Erin Kollins, Russell Byrum and Tracey Burdette for technical assistance with hamster experiments. This work was supported by the National Institute of Allergy and Infectious Diseases, National Institutes of Health, and by the Bill & Melinda Gates Foundation OPP1170236 and INV-004923 (I.A.W.). This work was also funded by the National Institutes of Health under R01AI132317 (D.N., D.H. and L.P.). This project has been funded in whole or in part with Federal funds from the National Institute of Allergy and Infectious Diseases, National Institutes of Health, Department of Health and Human Services, under Contract No. HHSN272201800013C.

## Author contributions

J.T. and P.D.C. conceived and designed the study. H.C., K.K.G-W., M.P. and J.T. isolated and characterized mAbs. D.H., J.L., E.P., C.A.T., W.S., E.S.Y., Y.Z., K.L., L.W., L.P., R.G., A.P. and M.H. performed SARS-CoV-2 binding and neutralization assays, as well as analysis of data. N.B., J.L. and Y.C. performed *in vivo* experiments. M.Y., J.L.T., S.B., N.C.W., and H.L. conducted structural work. J.S. performed statistics. S.L. and M.P. managed COVID-19 biospecimens. C.D. performed bead-based screening of antibodies. T.M., M.C. and I.D. performed cell sorting of COVID-19 samples. M.Z. performed liquid chromatography and purified antibodies. F.E-H.L. developed and provided plasmablast medium. R.S.W. organized the collection of COVID-19 samples. C.S., J.R.M., T.F.R., D.N., A.B.W., I.A.W., P.D.C. and J.T. aided in analysis and provided supervision.

