## Supplementary Figures for "Ultrapotent bispecific antibodies neutralize emerging SARS-CoV-2 variants"

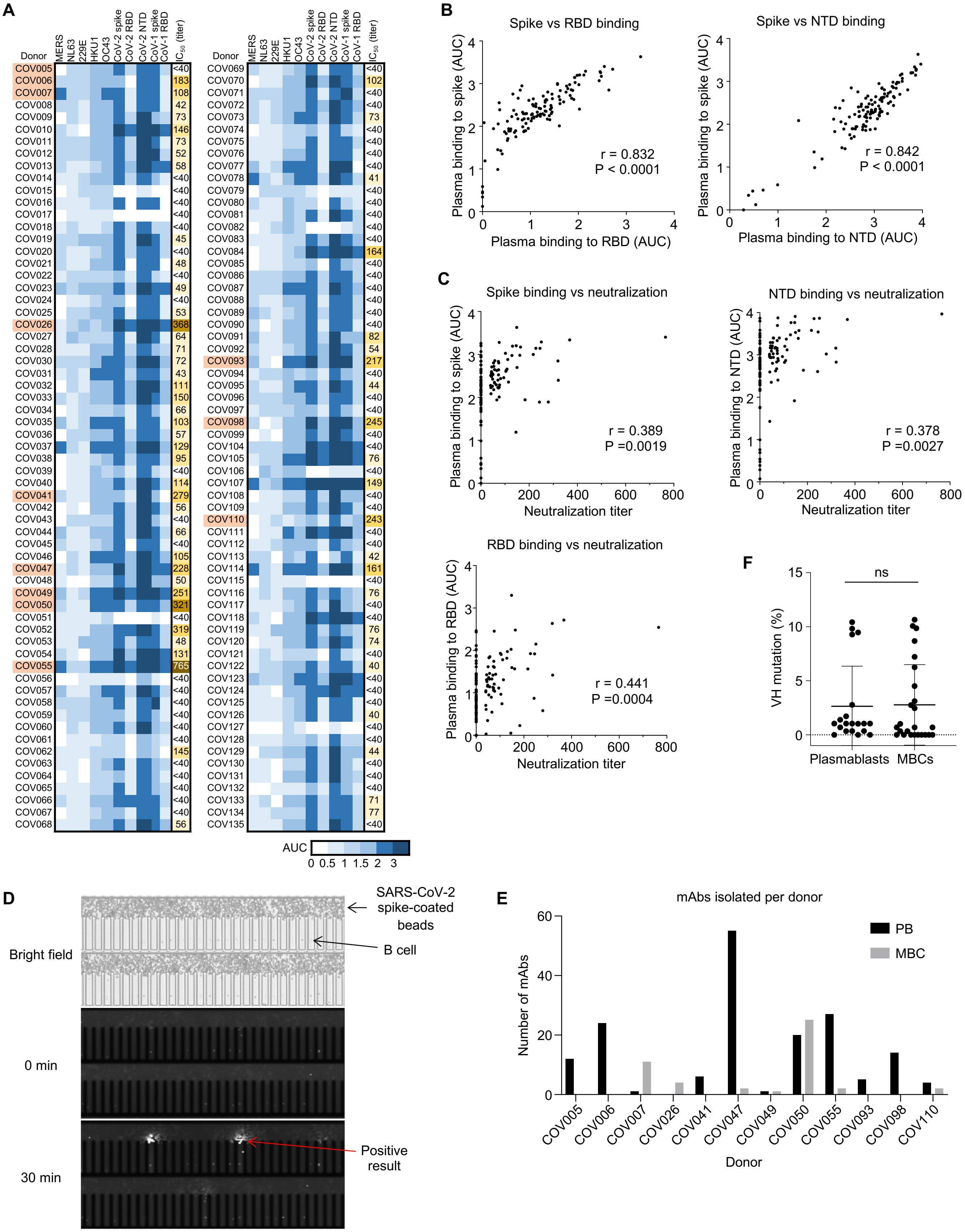
A
Donor
MERS
NL63
229E
HKU1
OC43
CoV-2 spike
CoV-2 RBD
CoV-2 NTD
CoV-1 spike
CoV-1 RBD
$IC_{50}$ (titer)
COV005 <40
COV006 183
COV007 108
COV008 42
COV009 73
COV010 146
COV011 73
COV012 52
COV013 58
COV014 <40
COV015 <40
COV016 <40
COV017 <40
COV018 <40
COV019 45
COV020 <40
COV021 48
COV022 <40
COV023 49
COV024 <40
COV025 53
COV026 368
COV027 64
COV028 71
COV030 72
COV031 43
COV032 111
COV033 150
COV034 66
COV035 103
COV036 57
COV037 129
COV038 95
COV039 <40
COV040 114
COV041 279
COV042 56
COV043 <40
COV044 66
COV045 <40
COV046 105
COV047 228
COV048 50
COV049 251
COV050 321
COV051 <40
COV052 319
COV053 48
COV054 131
COV055 765
COV056 <40
COV057 <40
COV058 <40
COV059 <40
COV060 <40
COV061 <40
COV062 145
COV063 <40
COV064 <40
COV065 <40
COV066 <40
COV067 <40
COV068 56
COV069 <40
COV070 102
COV071 <40
COV072 <40
COV073 73
COV074 <40
COV075 <40
COV076 <40
COV077 <40
COV078 41
COV079 <40
COV080 <40
COV081 <40
COV082 <40
COV083 <40
COV084 164
COV085 <40
COV086 <40
COV087 <40
COV088 <40
COV089 <40
COV090 <40
COV091 82
COV092 54
COV093 217
COV094 <40
COV095 44
COV096 <40
COV097 <40
COV098 245
COV099 <40
COV104 <40
COV105 76
COV106 <40
COV107 149
COV108 <40
COV109 <40
COV110 243
COV111 <40
COV112 <40
COV113 42
COV114 161
COV115 <40
COV116 76
COV117 <40
COV118 <40
COV119 76
COV120 74
COV121 <40
COV122 40
COV123 <40
COV124 <40
COV125 <40
COV126 40
COV127 <40
COV128 <40
COV129 44
COV130 <40
COV131 <40
COV132 <40
COV133 71
COV134 77
COV135 <40
AUC
0 0.5 1 1.5 2 3
B
Spike vs RBD binding
Plasma binding to spike (AUC)
Plasma binding to RBD (AUC)
r = 0.832
P < 0.0001
Spike vs NTD binding
Plasma binding to spike (AUC)
Plasma binding to NTD (AUC)
r = 0.842
P < 0.0001
C
Spike binding vs neutralization
Plasma binding to spike (AUC)
Neutralization titer
r = 0.389
P =0.0019
NTD binding vs neutralization
Plasma binding to NTD (AUC)
Neutralization titer
r = 0.378
P =0.0027
RBD binding vs neutralization
Plasma binding to RBD (AUC)
Neutralization titer
r = 0.441
P =0.0004
F
VH mutation (%)
ns
Plasmablasts
MBCs
D
SARS-CoV-2 spike-coated beads
B cell
Bright field
0 min
30 min
Positive result
E
mAbs isolated per donor
Number of mAbs
PB
MBC
COV005
COV006
COV007
COV026
COV041
COV047
COV049
COV050
COV055
COV093
COV098
COV110
Donor

**Supp. Fig. 1. Characterization of plasma and mAbs from COVID-19 convalescent donors. A**, Neutralization titers and values of plasma binding to the spike protein of multiple coronaviruses and specific domains of SARS-CoV-1 and SARS-CoV-2 (n = 126 donors). Area under the curve (AUC) values are shown after subtraction of the negative control antigen. Donors marked in orange were selected for mAb isolation. **B**, Associations between plasma binding to SARS-CoV-2 spike, RBD and NTD (n = 126 donors). P and r determined by Spearman's rank correlation. **C,** Relationship between neutralization and binding to SARS-CoV-2 spike, RBD and NTD (n = 126 donors). P and r determined by Spearman's rank correlation, plasma that were non-neutralizing at the highest concentration (1/40) are shown on the y-axis and were excluded from correlation analysis. **D,** Image of nanopens containing B cells. The lower panel shows positive fluorescence signals from binding of secreted antibodies to SARS-CoV-2 spike-coated beads. **E,** Number of mAbs isolated per donor, divided by cell type. **F,** VH mutations in mAbs isolated from donor COV050.

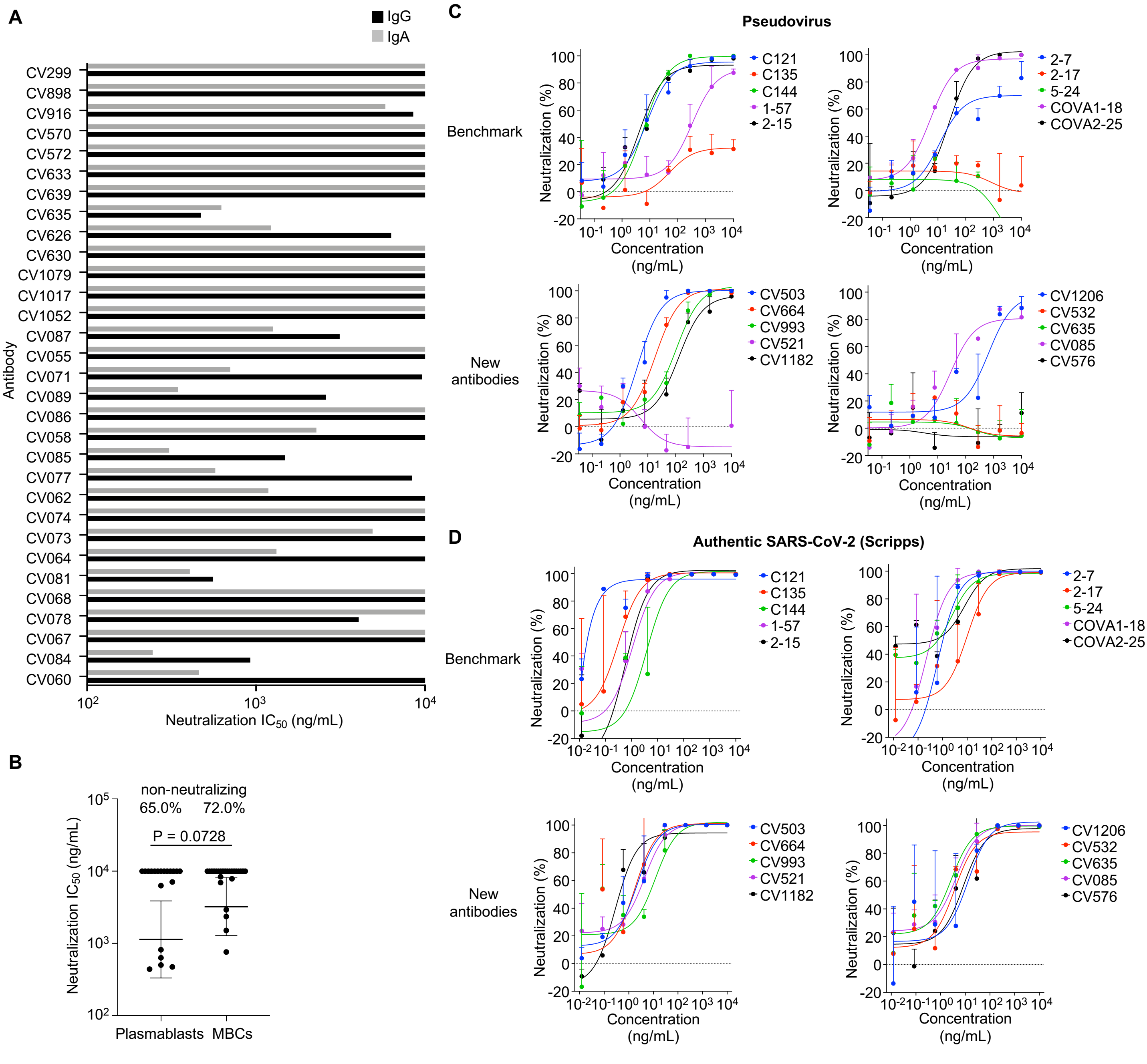


**Supp. Fig. 2. Pseudovirus and SARS-CoV-2 neutralization of benchmark and new antibodies. A**, Authentic SARS-CoV-2 neutralization (FRNA) potency of IgG and IgA forms of the same mAbs (originally isolated as IgA). **B**, Neutralization potency of antibodies isolated from donor COV050 by cell type. Top values indicate percentages of non-neutralizing antibodies. Bars indicate mean values ± SD; Mann-Whitney U-test (non-neutralizing antibodies excluded from calculation). **C,** Pseudovirus neutralization of antibody panel. Error bars show standard deviation. Representative of n = 3 experiments. **D**, Authentic SARS-CoV-2 neutralization (Scripps) of antibody panel. Error bars show standard deviation. Representative of n = 2 experiments.

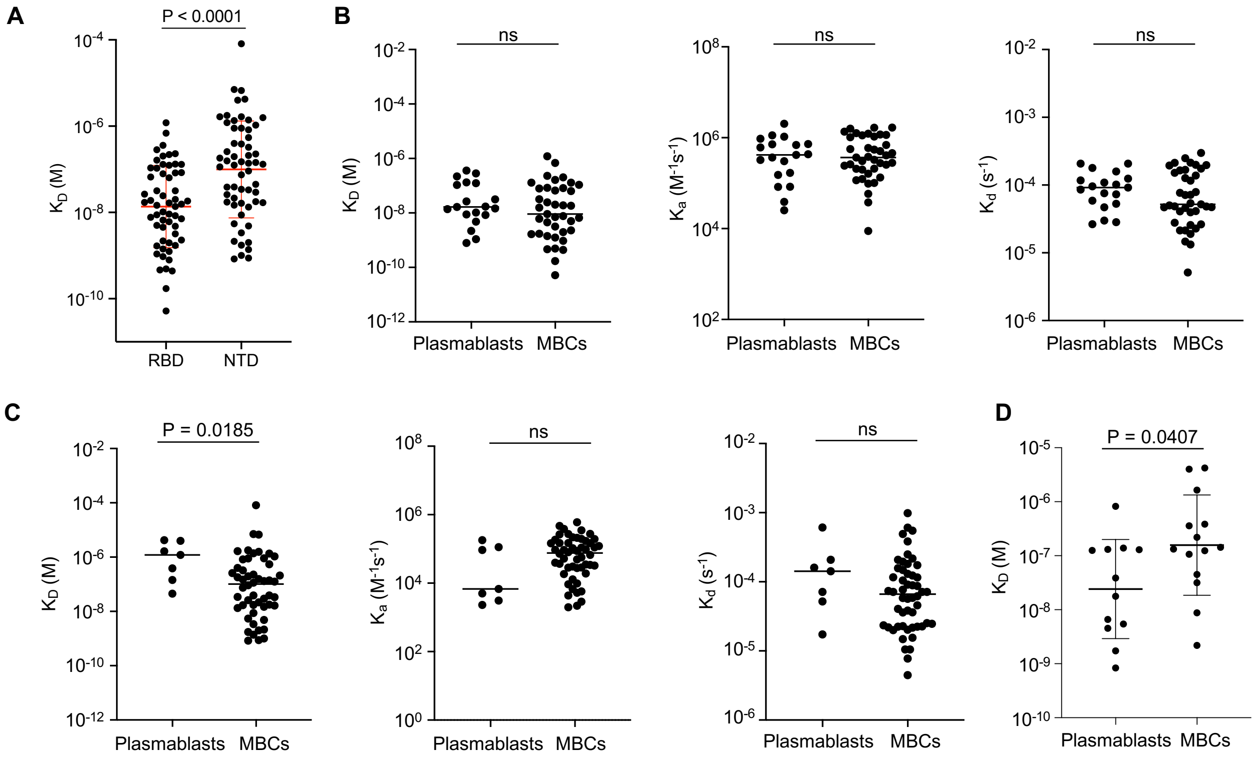


**Supp. Fig. 3. Comparison of affinity of antibodies from plasmablasts and memory B cells (MBCs). A**, Comparison of affinity of SARS-CoV-2 RBD-specific and NTD-specific antibodies. **B,** Comparison of affinity, association rates and dissociation rates of SARS-CoV-2 RBD-specific antibodies. **C,** Comparison of affinity, association rates and dissociation rates of SARS-CoV-2 NTD-specific antibodies. **D,** Comparison of affinity of antibodies from plasmablasts and MBCs isolated from donor COV050.

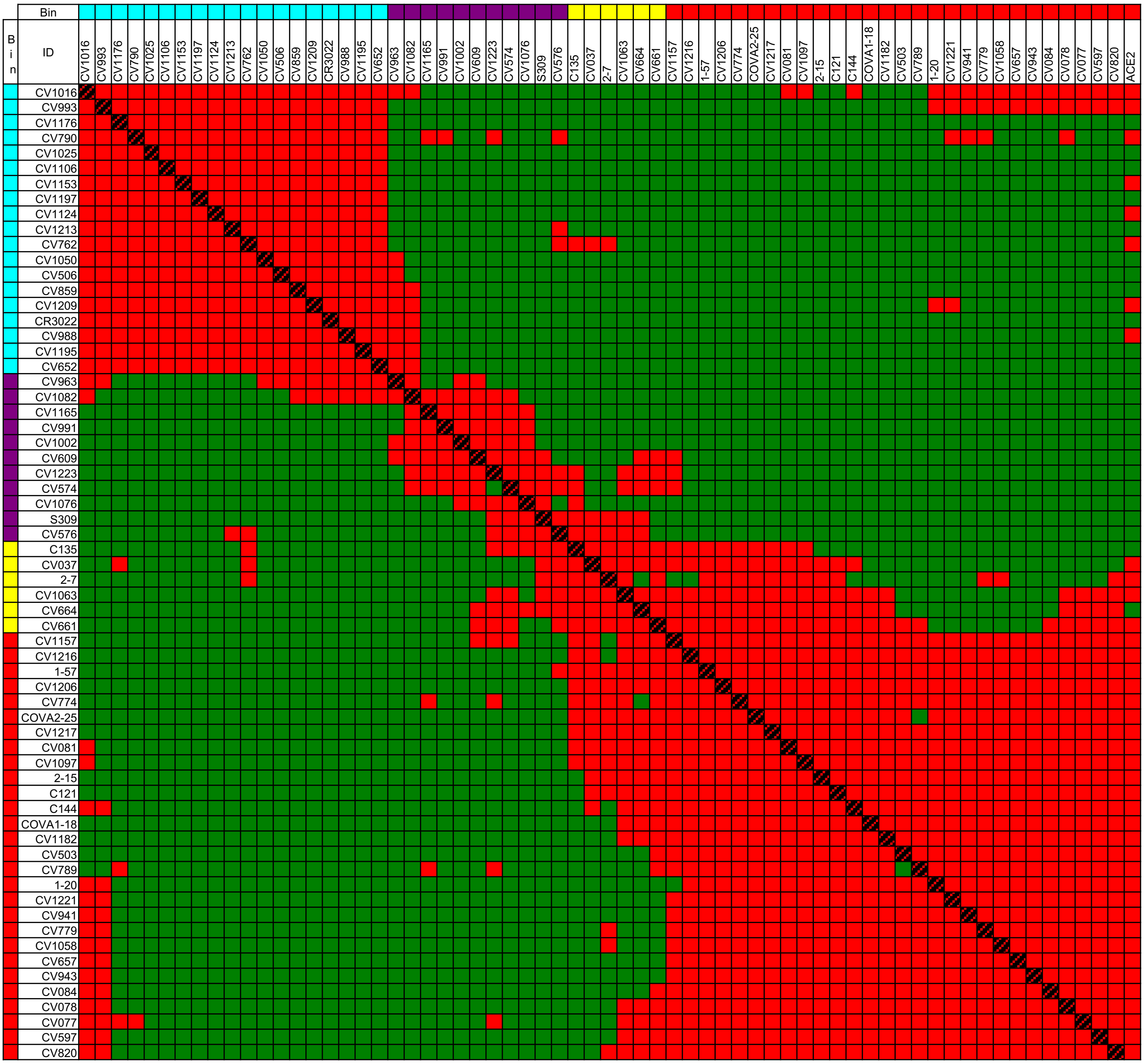


**Supp. Fig. 4. RBD epitope binning heat map.** A red square indicates competition while a green square indicates non-overlapping binding. The rows show antibodies attached to the SPR chip (ligands) while the columns show antibodies added after antigen binding (analytes). Representative of n = 2 experiments.

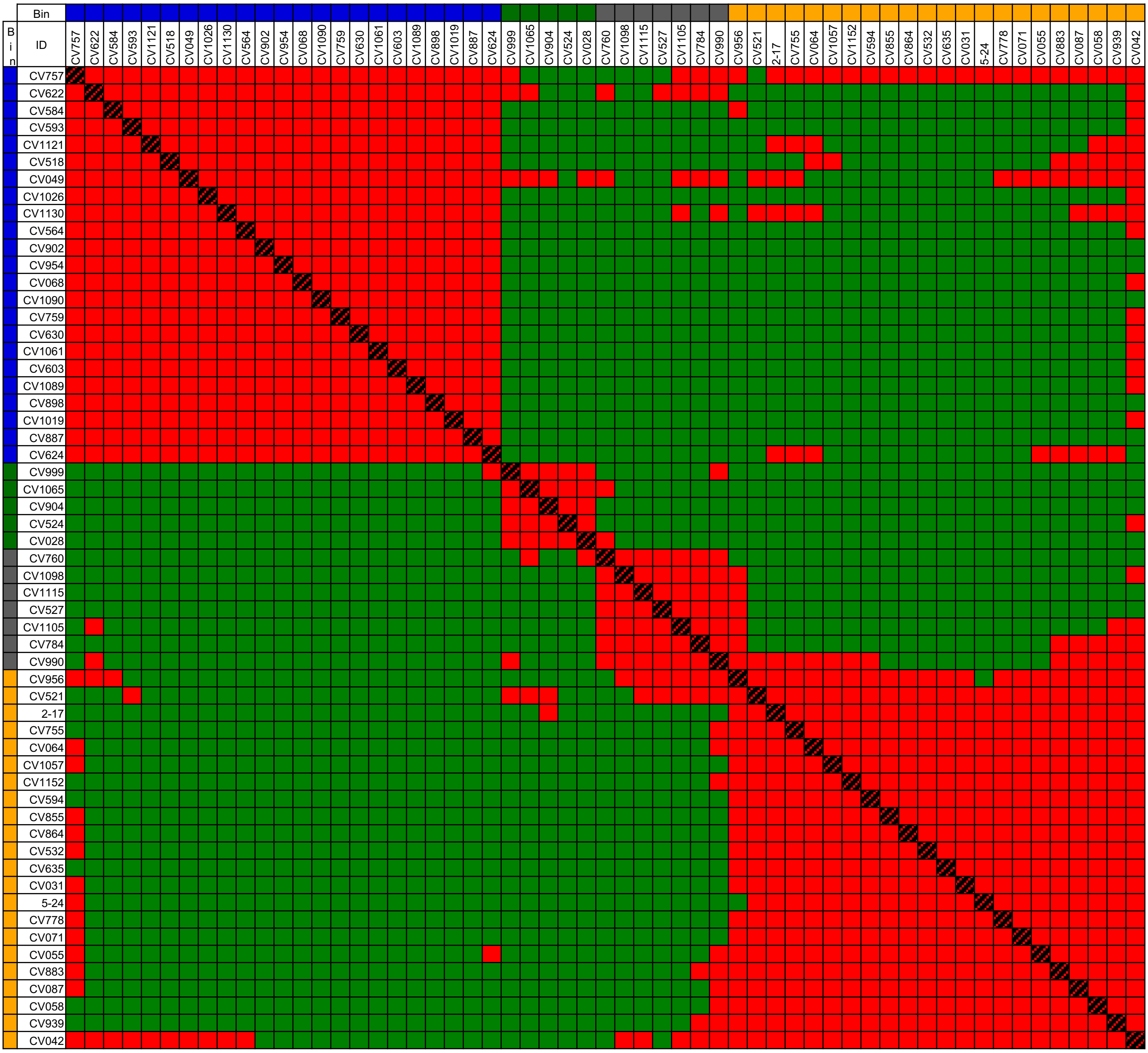


**Supp. Fig. 5. NTD epitope binning heat map.** A red square indicates competition while a green square indicates non-overlapping binding. The rows show antibodies attached to the SPR chip (ligands) while the columns show antibodies added after antigen binding (analytes). Representative of n = 2 experiments.

**A**

RBD-specific mAbs

| Antibody | D614G | B.1.1.7 | B.1.351 |
|---|---|---|---|
| CV037 | 100 | 133 | 155 |
| CV043 | 100 | 198 | 0 |
| CV047 | 100 | 123 | 236 |
| CV077 | 100 | 353 | -1 |
| CV078 | 100 | 138 | 0 |
| CV081 | 100 | 35 | 1 |
| CV084 | 100 | 290 | 8 |
| CV085 | 100 | 93 | 0 |
| CV089 | 100 | 15 | 0 |
| CV503 | 100 | 309 | 171 |
| CV506 | 100 | 215 | 136 |
| CV576 | 100 | 113 | 178 |
| CV597 | 100 | 196 | 2 |
| CV602 | 100 | 124 | 154 |
| CV626 | 100 | 110 | 0 |
| CV657 | 100 | 211 | 3 |
| CV661 | 100 | 104 | 1 |
| CV664 | 100 | 345 | 280 |
| CV774 | 100 | 594 | -2 |
| CV779 | 100 | -1 | -1 |
| CV789 | 100 | 120 | 114 |
| CV796 | 100 | 50 | 53 |
| CV820 | 100 | 0 | 0 |
| CV916 | 100 | 85 | 2 |
| CV941 | 100 | 174 | 0 |
| CV943 | 100 | 67 | 0 |
| CV958 | 100 | 120 | 1 |
| CV993 | 100 | 123 | 108 |
| CV1058 | 100 | 0 | 0 |
| CV1063 | 100 | 541 | 4 |
| CV1097 | 100 | 1 | 0 |
| CV1106 | 100 | 264 | 141 |
| CV1157 | 100 | 126 | 0 |
| CV1182 | 100 | 143 | 1 |
| CV1206 | 100 | 206 | 0 |
| CV1216 | 100 | 99 | 0 |
| CV1217 | 100 | 120 | 0 |

NTD-specific mAbs

| Antibody | D614G | B.1.1.7 | B.1.351 |
|---|---|---|---|
| CV031 | 100 | 0 | 0 |
| CV042 | 100 | 20 | 1 |
| CV071 | 100 | 3 | 2 |
| CV087 | 100 | 72 | 0 |
| CV280 | 100 | 12 | 3 |
| CV521 | 100 | 4 | -1 |
| CV524 | 100 | 137 | 116 |
| CV532 | 100 | 0 | 0 |
| CV584 | 100 | 121 | 60 |
| CV594 | 100 | 2 | 0 |
| CV635 | 100 | 29 | 0 |
| CV755 | 100 | 6 | 0 |
| CV778 | 100 | 0 | 3 |
| CV855 | 100 | 48 | 0 |
| CV864 | 100 | 0 | 0 |
| CV883 | 100 | 69 | 1 |
| CV956 | 100 | 102 | 153 |
| CV964 | 100 | 63 | 0 |
| CV1057 | 100 | 0 | 0 |
| CV1152 | 100 | 0 | 4 |

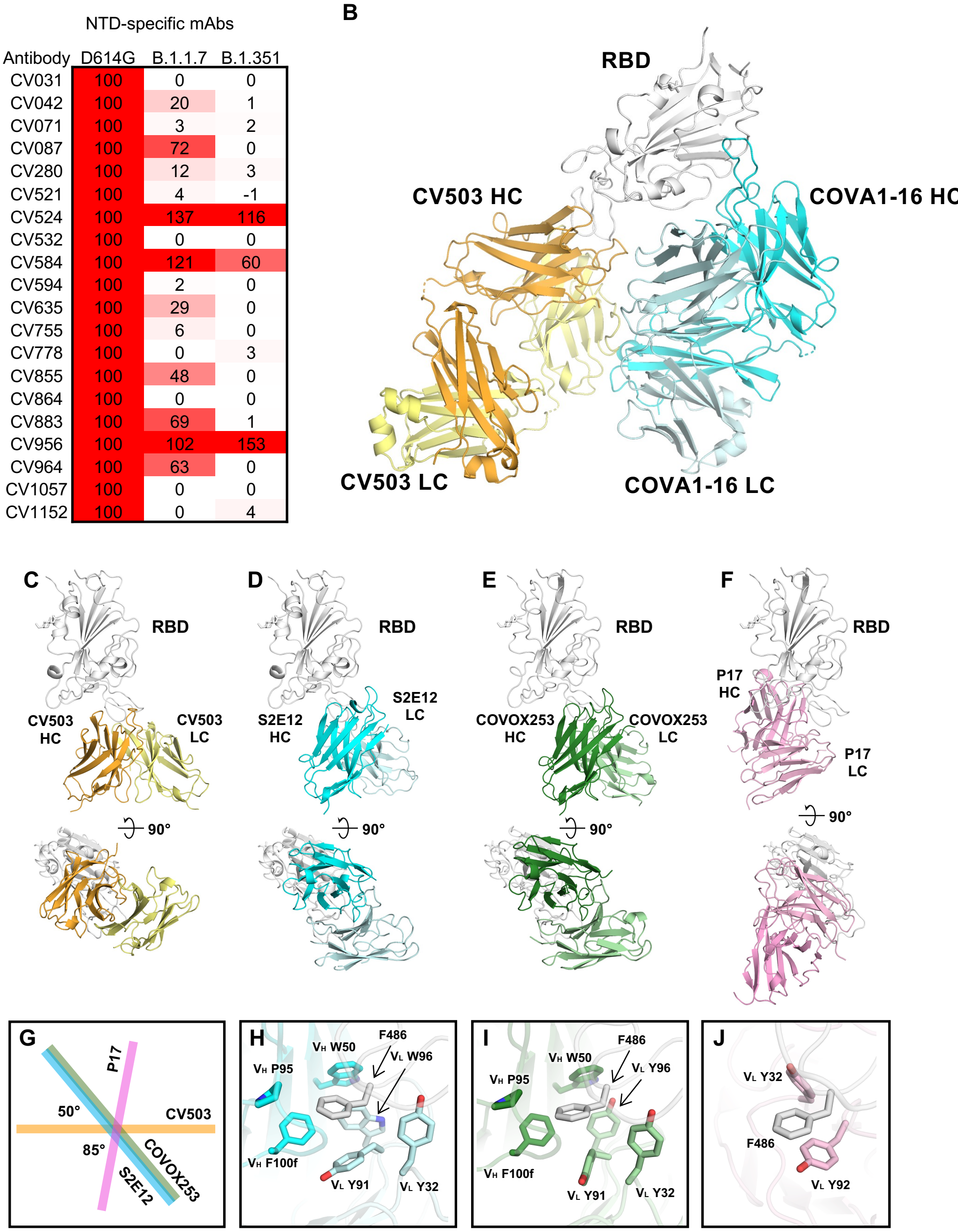


**Supp. Fig. 6. Binding of mAbs to SARS-CoV-2 spike protein. A,** Binding of all wild-type SARS-CoV-2-neutralizing antibodies (based on FRNA) to spike protein containing mutations from B.1.1.7 and B.1.351 variants (n = 1 experiment). The numbers show the percentages of mAb binding to mutants relative to D614G (which was normalized to 100). **B,** Crystal structure of SARS-CoV-2 RBD in complex with Fabs CV503 and COVA1-16. The binding site of CV503 with the Fab heavy and light chains shown in orange and yellow, respectively, on the RBD (white). The epitope is distinct from that of COVA1-16, whose Fab heavy and light chains are shown in cyan and pale cyan, respectively. **C-F,** Structures of CV503/RBD, S2E12/RBD (PDB ID: 7K45, ref. 20), COVOX-253/RBD (PDB ID: 7BEN, ref. 35), and P17/RBD (PDB ID: 7CWO, ref. 36) complexes are shown, where the RBD molecules (white) are shown in the same view. Only the variable domains of the Fabs are shown for clarity. **G,** S2E12 (cyan) and COVOX-253 (green) bind to RBD in a nearly identical approach, which are rotated by 50° compared to CV503 (orange), whereas P17 (pink) is rotated by 85° compared to CV503. The rotation angles are represented by insert. **H-I,** F486 of SARS-CoV-2 RBD (white) is clamped in a hydrophobic pocket formed by six aromatic residues from the heavy (cyan) and light chains (light cyan) of S2E12 **(H)**. A similar arrangement is seen for COVOX253 **(I)**. **J,** RBD-F486 stacks with $V_L$ 32 and $V_L$ 92 of P17 (pink). Kabat numbering is assigned to the antibody residues.

**A**

**Dose 1 (2x $IC_{50}$)**

| | Buffer | CV503 | CV521 | CV664 | CV993 | CV1182 |
|---|---|---|---|---|---|---|
| CV503 | 76.6 | | | | | |
| CV521 | 78.5 | 95.3 (95.0) | | | | |
| CV664 | 72.1 | 99.7 (93.5) | 92.8 (94.0) | | | |
| CV993 | 63.4 | 84.1 (91.4) | 95.0 (92.1) | 97.9 (89.8) | | |
| CV1182 | 69.7 | | 94.4 (93.5) | 81.9 (91.5) | 87.8 (88.9) | |

**Dose 2 (1x $IC_{50}$)**

| | Buffer | CV503 | CV521 | CV664 | CV993 | CV1182 |
|---|---|---|---|---|---|---|
| CV503 | 40.0 | | | | | |
| CV521 | 61.3 | 85.3 (76.7) | | | | |
| CV664 | 39.7 | 85.0 (63.8) | 78.1 (76.6) | | | |
| CV993 | 39.3 | 64.5 (63.6) | 79.1 (76.5) | 78.8 (63.4) | | |
| CV1182 | 36.5 | | 74.8 (75.4) | 71.0 (61.7) | 69.7 (61.5) | |

**B**

**CV503 + CV664**

**Run 1**

| | | CV664 (ng/mL) | | | | | |
|---|---|---|---|---|---|---|---|
| | | 0.0 | 19.8 | 39.6 | 79.2 | 158.3 | 316.7 |
| CV503 (ng/mL) | 0.0 | 0.0 | 15.0 | -4.9 | 23.0 | 25.3 | 45.9 |
| | 14.8 | 24.9 | 12.6 | 19.3 | 27.5 | 19.5 | 55.4 |
| | 29.6 | 14.4 | 11.2 | 27.2 | 25.1 | 47.0 | 54.7 |
| | 59.2 | 13.2 | 17.2 | 32.1 | 36.5 | 46.6 | 70.5 |
| | 118.4 | 41.9 | 36.5 | 47.6 | 68.6 | 72.1 | 91.8 |
| | 236.8 | 55.0 | 56.2 | 59.9 | 64.2 | 74.0 | 97.4 |

**Run 2**

| | | CV664 (ng/mL) | | | | | |
|---|---|---|---|---|---|---|---|
| | | 0.0 | 19.8 | 39.6 | 79.2 | 158.3 | 316.7 |
| CV503 (ng/mL) | 0.0 | 0.0 | 8.5 | 20.7 | 29.8 | 51.6 | 63.1 |
| | 14.8 | 28.9 | 25.0 | 36.0 | 36.9 | 29.7 | 84.3 |
| | 29.6 | 0.6 | 21.1 | 47.1 | 54.7 | 43.1 | 91.0 |
| | 59.2 | -2.3 | 21.8 | 61.3 | 70.2 | 80.6 | 98.1 |
| | 118.4 | 25.4 | 21.1 | 68.8 | 83.6 | 97.3 | 99.3 |
| | 236.8 | 55.2 | 68.2 | 92.9 | 92.6 | 99.1 | 99.4 |

Synergy values (Loewe's, >10 indicates synergy)
Run 1: 6.99
Run 2: 18.23

**CV664 + CV993**

**Run 1**

| | | CV664 (ng/mL) | | | | | |
|---|---|---|---|---|---|---|---|
| | | 0.0 | 19.8 | 39.6 | 79.2 | 158.3 | 316.7 |
| CV993 (ng/mL) | 0.0 | 0.0 | 15.0 | -4.9 | 23.0 | 25.3 | 45.9 |
| | 11.6 | 8.9 | 9.6 | 18.4 | 28.2 | 34.3 | 46.2 |
| | 23.2 | 15.7 | 20.5 | 25.5 | 20.3 | 18.7 | 68.8 |
| | 46.5 | 20.5 | 22.3 | 32.7 | 18.9 | 43.2 | 66.5 |
| | 92.9 | 34.5 | 32.9 | 28.9 | 30.7 | 61.5 | 66.4 |
| | 185.8 | 45.4 | 49.9 | 59.4 | 55.9 | 75.7 | 90.9 |

**Run 2**

| | | CV664 (ng/mL) | | | | | |
|---|---|---|---|---|---|---|---|
| | | 0.0 | 19.8 | 39.6 | 79.2 | 158.3 | 316.7 |
| CV993 (ng/mL) | 0.0 | 0.0 | 8.5 | 20.7 | 29.8 | 51.6 | 63.1 |
| | 11.6 | 21.0 | 14.5 | 1.6 | 35.5 | 30.2 | 90.7 |
| | 23.2 | 22.9 | 29.7 | 20.6 | 51.0 | 22.6 | 86.6 |
| | 46.5 | 13.0 | 33.8 | 37.9 | 37.8 | 53.6 | 90.5 |
| | 92.9 | 25.6 | 37.5 | 58.0 | 76.2 | 88.1 | 96.9 |
| | 185.8 | 35.5 | 44.5 | 69.1 | 74.2 | 98.3 | 99.0 |

Synergy values (Loewe's, >10 indicates synergy)
Run 1: 3.81
Run 2: 19.31

**C**

**CV503 + CV664**

| | | CV664 (ng/mL) | | | | | | | |
|---|---|---|---|---|---|---|---|---|---|
| | | 0.0 | 0.1 | 0.4 | 1.2 | 3.7 | 11.1 | 33.3 | 100.0 |
| CV503 (ng/mL) | 0.0 | 37.5 | 28.4 | -8.9 | -5.1 | 77.8 | 89.8 | 99.4 | 100.0 |
| | 0.1 | 71.4 | 23.0 | 66.6 | 8.6 | 90.5 | 72.1 | 99.6 | 100.0 |
| | 0.4 | 27.0 | 40.8 | 67.3 | 24.5 | 83.1 | 94.3 | 99.6 | 100.0 |
| | 1.2 | 67.4 | 44.4 | 45.9 | 88.5 | 73.5 | 97.8 | 100.0 | 99.1 |
| | 3.7 | 83.3 | 75.1 | 94.3 | 40.4 | 94.3 | 99.6 | 100.0 | 100.0 |
| | 11.1 | 94.7 | 98.1 | 80.2 | 98.4 | 99.8 | 100.0 | 100.0 | 100.0 |
| | 33.3 | 100.0 | 98.7 | 99.8 | 99.8 | 100.0 | 100.0 | 100.0 | 100.0 |
| | 100.0 | 100.0 | 100.0 | 100.0 | 100.0 | 100.0 | 100.0 | 100.0 | 100.0 |

**CV664 + CV993**

| | | CV993 (ng/mL) | | | | | | | |
|---|---|---|---|---|---|---|---|---|---|
| | | 0.0 | 0.1 | 0.4 | 1.2 | 3.7 | 11.1 | 33.3 | 100.0 |
| CV664 (ng/mL) | 0.0 | 48.2 | 37.1 | -11.2 | 37.2 | -11.4 | 86.4 | 68.3 | 98.7 |
| | 0.1 | 42.7 | -20.6 | 30.1 | 40.9 | 41.3 | 79.3 | 100.0 | 99.6 |
| | 0.4 | -36.3 | 28.3 | -2.1 | 41.8 | 56.5 | 62.5 | 93.6 | 100.0 |
| | 1.2 | 37.3 | 19.8 | -12.4 | 16.8 | 44.1 | 78.8 | 100.0 | 99.7 |
| | 3.7 | 24.5 | 30.6 | 76.6 | -11.6 | 93.4 | 100.0 | 100.0 | 100.0 |
| | 11.1 | 100.0 | 96.7 | 67.2 | 97.5 | 86.5 | 100.0 | 100.0 | 100.0 |
| | 33.3 | 100.0 | 98.4 | 100.0 | 100.0 | 100.0 | 100.0 | 100.0 | 100.0 |
| | 100.0 | 100.0 | 100.0 | 100.0 | 100.0 | 100.0 | 100.0 | 100.0 | 100.0 |

Synergy values (Loewe's, >10 indicates synergy)
CV503 + CV664: -3.88
CV664 + CV993: 9.08

**Supp. Fig. 7. Screening of antibody combinations for synergy in neutralizing SARS-CoV-2. A**, Screen of antibody combinations for synergy in neutralizing authentic SARS-CoV-2 at two different antibody doses (2 x $IC_{50}$ and $IC_{50}$). Only non-overlapping pairs based on epitope binning data were tested. The numbers outside the brackets show the observed neutralization percentages while numbers in the brackets show the expected values. Combinations with an observed neutralization percentage of >5% the expected value are highlighted blue. **B**, Neutralization of authentic SARS-CoV-2 (FRNA assay) by titrations of CV503 and CV664, as well as CV664 and CV993. The numbers in the heat map show the neutralization percentages, and the numbers at the side show synergy scores for each run. **C**, Neutralization of authentic SARS-CoV-2 (Scripps assay) by titrations of CV503 and CV664, as well as CV664 and CV993. The numbers in the heat map show the neutralization percentages, and the numbers at the side show synergy scores for each run.

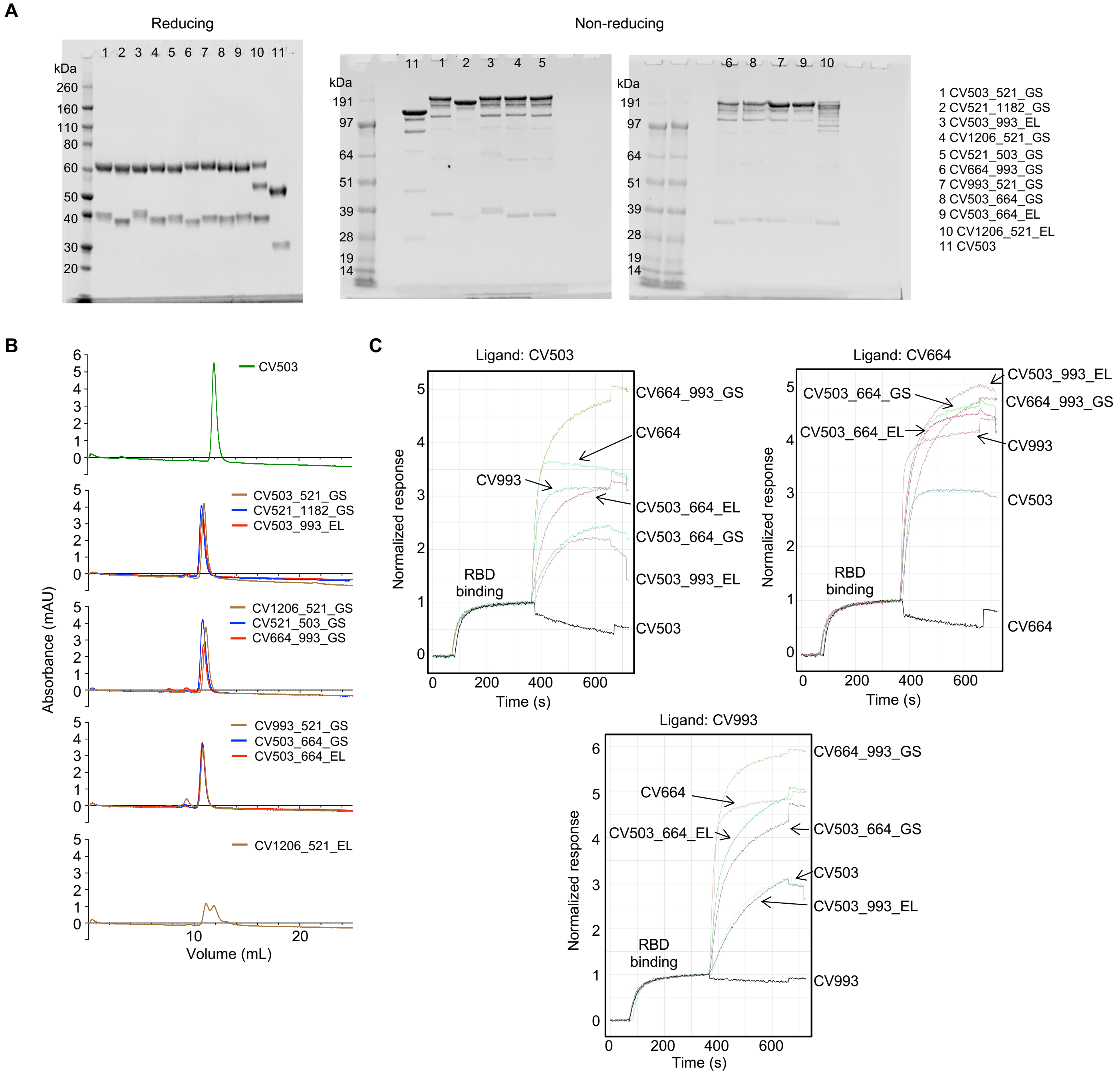


**Supp. Fig. 8. Bispecific antibodies are expressed as a single dominant species and use both binding sites. A,** Molecular weight of bispecific antibodies by SDS-PAGE. CV503 is a control standard IgG. In our bispecific antibody naming system, the first name refers to the antibody used to make the outer binding site and the second refers to the antibody at the inner binding site. GS or EL refers to the type of linker connecting the two antigen-binding sites. See Materials and Methods for details. **B**, Size exclusion chromatography of bispecific antibodies, compared to the control standard IgG CV503. **C**, Sensograms showing binding of bispecific antibodies to RBD attached to ligand IgGs (which are conjugated to the SPR chip). An increase in signal after RBD binding indicates attachment of the second antibody. Representative of n = 2 experiments.

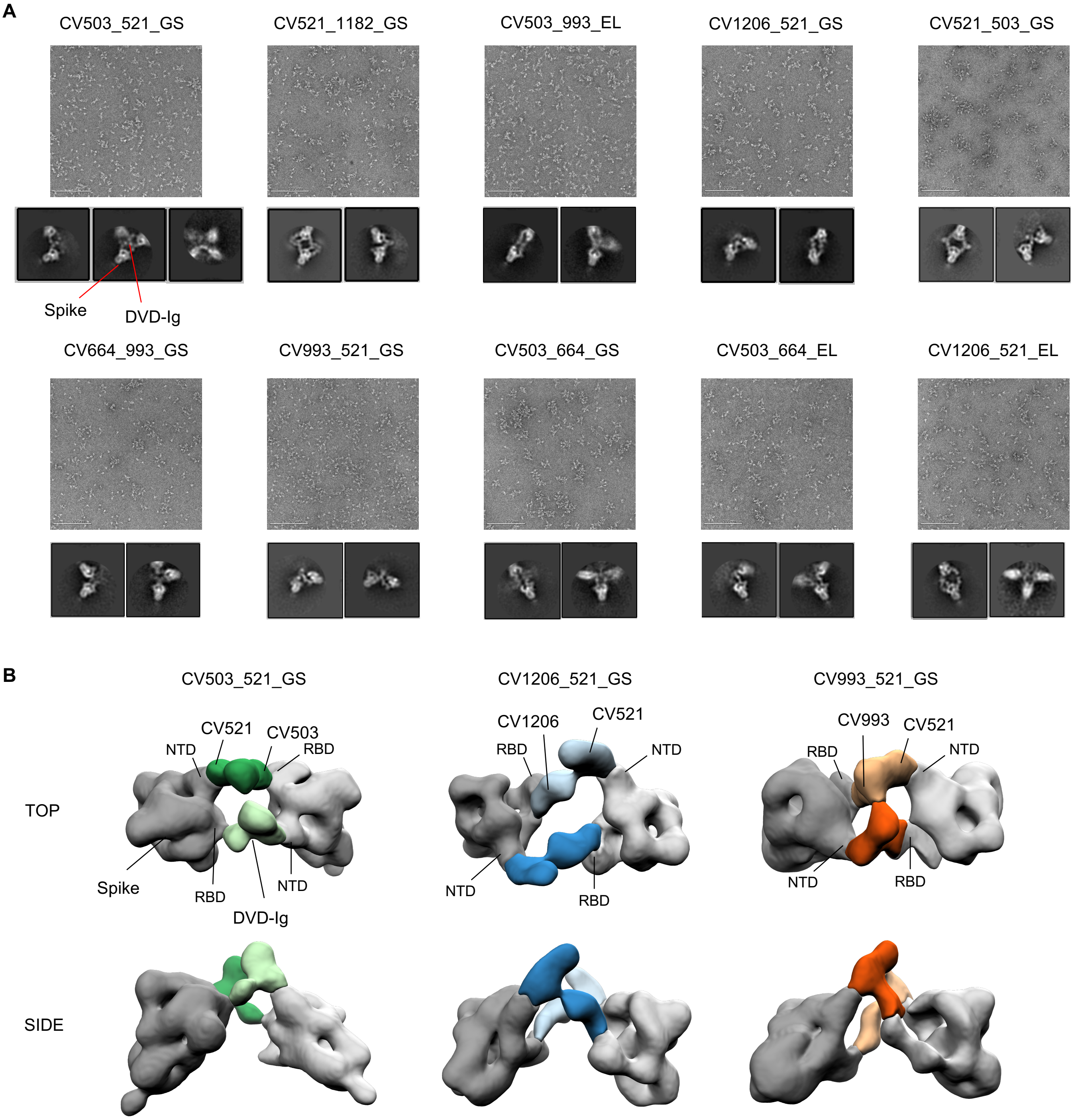


**Supp. Fig. 9. Negative stain EM images of bispecific antibodies in complex with spike protein. A**, Raw micrograph exemplars and select 2D classes of the bispecific antibody panel in complex with SARS-2 CoV 6P Mut7 spike protein. 2D classes show the DVD-Ig induced crosslinking of 2-4 spike proteins. **B**, Segmented 3D refinements from the negative stain EM data. Only 3 out of the 10 bispecific antibodies in complex with SARS-2 CoV 6P Mut7 were able to converge in 3D. Spike proteins are in gray and the bispecific antibodies are colored in green, blue, and orange for CV503_521_GS, CV1206_521_GS, and CV993_521_GS, respectively.

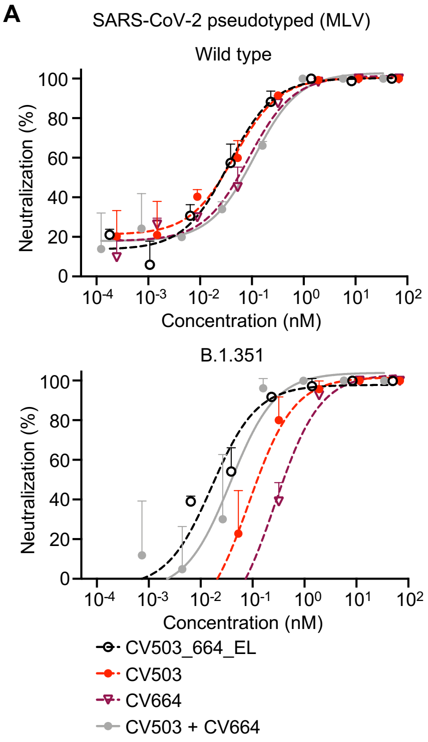


**B**

| Parameter | | Score |
|---|---|---|
| **Natural behavior** | **-normal** | **0** |
| | **-lethargy or staggered gait** | **1** |
| | **-Increased respiration, decreased mobility** | **3** |
| | **-greatly labored (decreased) respiration, gasping, or inability to move** | **5** |
| **Provoked behavior** | **-normal** | **0** |
| | **-subdued but normal when stimulated** | **1** |
| | **-subdued even when stimulated** | **3** |
| | **-unresponsive when stimulated** | **5** |
| **Total score** | **0 normal** | |
| | **1-2 normal observations (1x/day)** | |
| | **3 monitor with increased frequency (2x day)** | |
| | **≥5 moribund – euthanize** | |

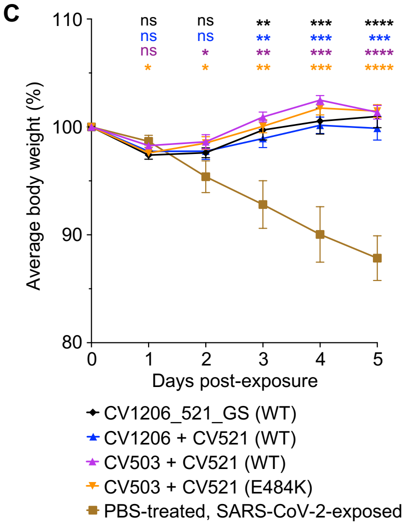


**Supp. Fig. 10. *In vitro* and *in vivo* potency of bispecific antibodies targeting SARS-CoV-2. A,** Potency of CV503_664_EL versus individual component mAbs against wild-type and B.1.351 SARS-CoV-2 pseudotyped virus (MLV). **B,** Clinical criteria used to evaluate individual hamsters exposed to SARS-CoV-2. **C,** Weight change in hamsters that were administered bispecific antibodies at 1 mg/hamster or an equimolar mAb cocktail (0.72-0.73 mg of each mAb/hamster), 12 h prior to IN virus exposure at 5log10 PFU. Differences between groups that were given the antibody versus PBS were determined using a mixed-effects repeated measures analysis with Dunnett's multiple comparisons; *P<0.05, **P<0.01, ***P<0.001, ****P<0.0001. n = 5 hamsters per group. Points represent mean +/- standard deviation (SD).
