## Supplementary Table 2 for "Ultrapotent bispecific antibodies neutralize emerging SARS-CoV-2 variants"

### Supplementary Table 2. X-ray data collection and refinement statistics

| **Data collection** | CV503 + RBD + COVA1-16 |
|---|---|
| Beamline | SSRL12-1 |
| Wavelength (Å) | 0.97946 |
| Space group | P 1 $2_1$ 1 |
| Unit cell parameters | |
| a, b, c (Å) | 172.2, 122.7, 175.5 |
| α, β, γ (°) | 90, 118.2, 90 |
| Resolution (Å) [a] | 50.0-3.40 (3.48-3.40) |
| Unique reflections [a] | 87,443 (8,562) |
| Redundancy [a] | 3.4 (3.5) |
| Completeness (%) [a] | 98.8 (99.8) |
| $<I/\sigma_I>$ [a] | 10.2 (1.0) |
| $R_{sym}$[b] (%) [a] | 14.6 (>100) |
| $R_{pim}$[b] (%) [a] | 4.3 (64.9) |
| $CC_{1/2}$[c] (%) [a] | 99.8 (53.6) |
| **Refinement statistics** | |
| Resolution (Å) | 40.6-3.40 |
| Reflections (work) | 87,416 |
| Reflections (test) | 2,000 |
| $R_{cryst}$[d] / $R_{free}$[e] (%) | 20.0/23.5 |
| No. of atoms | 24,307 |
| RBD | 4,671 |
| CV503 Fab | 9,602 |
| COVA1-16 Fab | 10,034 |
| Average *B*-values (Å$^2$) | 131 |
| RBD | 139 |
| CV503 Fab | 131 |
| COVA1-16 Fab | 128 |
| Wilson *B*-value (Å$^2$) | 128 |
| **RMSD from ideal geometry** | |
| Bond length (Å) | 0.002 |
| Bond angle (°) | 0.60 |
| **Ramachandran statistics (%)** | |
| Favored | 95.8 |
| Outliers | 0.28 |
| **PDB code** | pending |

[a] Numbers in parentheses refer to the highest resolution shell.

[b] $R_{sym} = \Sigma_{hkl} \Sigma_i | I_{hkl,i} - <I_{hkl}> | / \Sigma_{hkl} \Sigma_i I_{hkl,i}$ and $R_{pim} = \Sigma_{hkl} (1/(n-1))^{1/2} \Sigma_i | I_{hkl,i} - <I_{hkl}> | / \Sigma_{hkl} \Sigma_i I_{hkl,i}$, where $I_{hkl,i}$ is the scaled intensity of the $i^{th}$ measurement of reflection h, k, l, $<I_{hkl}>$ is the average intensity for that reflection, and *n* is the redundancy.

[c] $CC_{1/2}$ = Pearson correlation coefficient between two random half datasets.

[d] $R_{cryst} = \Sigma_{hkl} | F_o - F_c | / \Sigma_{hkl} | F_o | \times 100$, where $F_o$ and $F_c$ are the observed and calculated structure factors, respectively.

[e] $R_{free}$ was calculated as for $R_{cryst}$, but on a test set comprising 5% of the data excluded from refinement.
